# The glycoprotein quality control factor Malectin promotes coronavirus replication and viral protein biogenesis

**DOI:** 10.1101/2024.06.02.597051

**Authors:** Jonathan P. Davies, Lars Plate

**Affiliations:** Department of Biological Sciences, Vanderbilt University, Nashville, TN, 37235; Vanderbilt Institute of Infection, Immunology and Inflammation, Nashville, TN, 37235; Department of Chemistry, Vanderbilt University, Nashville, TN, 37235; Department of Pathology, Microbiology and Immunology, Vanderbilt University Medical Center, Nashville, TN, 37235

**Keywords:** Coronavirus, malectin, affinity-purification mass spectrometry (AP-MS), oligosaccharyltransferase (OST) complex

## Abstract

Coronaviruses (CoV) rewire host protein homeostasis (proteostasis) networks through interactions between viral nonstructural proteins (nsps) and host factors to promote infection. With the emergence of SARS-CoV-2, it is imperative to characterize host interactors shared across nsp homologs. Using quantitative proteomics and functional genetic screening, we identify conserved proteostasis interactors of nsp2 and nsp4 that serve pro-viral roles during infection of murine hepatitis virus – a model betacoronavirus. We uncover a glycoprotein quality control factor, Malectin (MLEC), which significantly reduces infectious titers when knocked down. During infection, nsp2 interacts with MLEC-associated proteins and the MLEC-interactome is drastically altered but retains association with the Oligosaccheryltransferase (OST) complex, a crucial component of viral glycoprotein production. MLEC promotes viral protein levels and genome replication through its quality control activity. Lastly, we show MLEC promotes SARS-CoV-2 replication. Our results reveal a role for MLEC in mediating CoV infection and identify a potential target for pan-CoV antivirals.

## Introduction

Several highly pathogenic human coronaviruses (CoVs) have emerged in the past few decades and imposed drastic public health burdens, including the betacoronaviruses SARS-CoV, MERS-CoV, and most recently SARS-CoV-2. Given the pattern of past CoV emergence and the likelihood of future outbreaks, it is imperative to understand the molecular strategies shared by CoVs to infect human cells. CoVs hijack host cellular systems through virus-host protein-protein interactions (PPIs) to mediate infection(Perlman and Netland, 2009). In particular, the sixteen nonstructural proteins (nsps) encoded by CoVs alter the cellular environment to enhance infection and create viral replication/transcription complexes (RTCs) to reproduce viral genomes(V’kovski et al., 2021).

Nsps have varying degrees of sequence conservation, topology, and ascribed functions. This work focuses on two nsps involved in setting up viral replication centers: nsp2 and nsp4. Both viral proteins are translated early in infection but have contrasting degrees of known functions. We previously characterized the comparative host interactomes of nsp2 and nsp4(Davies et al., 2020) and herein describe our functional interrogation of these interactions.

Nsp2 is a soluble protein with an ambiguous role in viral infection and low sequence conservation amongst CoV homologs. Previous work has found that nsp2 can be deleted from the Murine Hepatitis Virus (MHV) and SARS-CoV genome and still produce infectious virus, albeit with significant stunting in titers (∼two log_10_ plaque forming units (PFU)/mL)(Graham et al., 2005). Nsp2 has also been shown to associate with other nsps amid the viral RTC(Freeman et al., 2014; Hagemeijer et al., 2009; Prentice et al., 2004b), indicating that while dispensable, nsp2 promotes viral infection and likely works in coordination with other viral proteins. Nsp4 is a multipass transmembrane glycoprotein with high sequence conservation across CoV homologs(Beachboard et al., 2015; Gadlage et al., 2010; Oostra et al., 2007). In conjunction with nsp3 and nsp6, nsp4 plays an indispensable role in host membrane manipulation to form double-membrane vesicles (DMVs), which serve as the primary sites of viral genome replication (Angelini et al., 2013; Oudshoorn et al., 2017; Sparks et al., 2007; Tabata et al., 2021).

One of the primary cellular systems hijacked by CoVs during infection is the host proteostasis network. The proteostasis network maintains the cellular proteome, from protein folding, addition of post-translational modifications, and trafficking to degrading old or misfolded proteins(Balch et al., 2008). Key to this network is the coordination of PPIs between quality control factors and client proteins to surveil the state of proteome, which is particularly important in the biogenesis and secretion of glycosylated proteins (glycoproteins)(Guay et al., 2023; Hammond et al., 1994; Wright et al., 2022, 2021). CoVs encode several *N*-linked glycoproteins with critical roles in replication, including nsp3, nsp4, spike (S), and membrane (M) proteins, while MHV encodes an additional structural glycoprotein, hemagglutinin-esterase (HE)(Ujike and Taguchi, 2015). Therefore, CoVs rely on host glycoprotein biogenesis machinery to produce viral glycoproteins.

Previous work has harnessed quantitative proteomics to characterize interactions between nsp2 and the host proteostasis system. V’kovski et al. applied proximity-labeling using a BioID-nsp2 encoding MHV strain to identify host factors proximal to the RTC, including autophagy and translation initiation factors(V’kovski et al., 2019). Nsp-host interactors of have also been identified using affinity-purification mass spectrometry (AP-MS) in multiple studies(Almasy et al., 2021b; Cornillez-Ty et al., 2009; Davies et al., 2020; David E. Gordon et al., 2020; David E Gordon et al., 2020; Stukalov et al., 2021). These studies have consistently identified the host translation repressor complex of 4EHP/GIGYF2 as an interactor of SARS-CoV and SARS-CoV-2 nsp2 homologs, and subsequent work has characterized the structural determinants of this interaction(Gupta et al., 2021; Xu et al., 2022; Zou et al., 2022). MERS-CoV nsp2 interacts with components of the Ribosome Quality Control (RQC) pathway(David E Gordon et al., 2020), including the ASC-1 complex(Juszkiewicz et al., 2020) and TCF25(Kuroha et al., 2018), in addition to a component of *O*-linked oligosaccharide biosynthesis (GALNT2)(Zilmer et al., 2020). However, the interface between nsp2 and glycoprotein biogenesis pathways has remained unexplored.

Interactions between nsp4 and endoplasmic reticulum (ER) proteostasis factors have also been examined, particularly in the context of DMV formation(Pahmeier et al., 2023). Various ER interactors have been shown to promote formation of DMVs from ER-derived membranes during SARS-CoV-2 infection, including RTN3/4(Williams et al., 2023), VMP1, and TMEM41B(Ji et al., 2022). SARS-CoV-2 nsp4 has also been found to activate the Unfolded Protein Response (UPR), a cellular stress response pathway, which mitigates the accumulation of misfolded proteins in part through ER expansion and ER chaperone upregulation(Davies et al., 2023).

Previously, we performed a comparative interactomics study of nsp2 and nsp4 homologs from SARS-CoV-2, SARS-CoV, and hCoV-OC43(Davies et al., 2020). We identified several conserved proteostasis interactors amongst SARS-CoV-2 and SARS-CoV nsp2 homologs, including the chaperones HSPA5 and HSPA8, the glycoprotein quality control factor MLEC(Galli et al., 2011; Schallus et al., 2008), and the ubiquitination complex ERLIN1/2/RNF170(Lu et al., 2011; Wright et al., 2015). We also found conserved nsp4-proteostasis interactors, such as components of the *N*-linked glycosylation machinery (DDOST, STT3B, MAGT1, CANX), the ER Membrane Complex (MMGT1)(Bai et al., 2020; Goytain and Quamme, 2008), ER-cargo receptors (CCPG1(Smith et al., 2018), SURF4(Emmer et al., 2018)), and protein degradation factors (TMEM33, TMEM43, HERPUD1, ERLIN1/2).

In this work, we expand our interactomics to a wider panel of nsp2 and nsp4 homologs, including MERS-CoV, hCoV-229E, and MHV. We evaluate the functional relevance of the PPIs and find that many conserved ER proteostasis interactors serve pro-viral roles in a CoV infection model, most notably the glycoprotein quality control factor Malectin (MLEC). We interrogate the nature of the interaction between nsp2 and find that MLEC aids viral protein production and genome replication during infection. Importantly, we find that MLEC promotes both MHV and SARS-CoV-2 replication. Our work identifies a new virus-host dependency with potential for pan-CoV therapeutics and furthers our understanding of the basic biology of CoV infection.

## Results

### Nsp2 and nsp4 ER proteostasis interactors are conserved across CoV homologs

We previously mapped the comparative interactome of nsp2 (SARS-CoV-2 and SARS-CoV) and nsp4 (SARS-CoV-2, SARS-CoV, hCoV-OC43) homologs(Davies et al., 2020). To determine the scope of conservation of previously identified interactors, we expanded our panel of nsp homologs to include the alphacoronavirus hCoV-229E and the betacoronaviruses MERS-CoV and Murine Hepatitis Virus (MHV) (**Fig. 1A**). The MHV homologs were included to determine the relevance of using MHV as a BSL-2 infection model for testing the functionality of interactors in active infection. We utilized our previously reported nsp construct design and affinity-purification mass spectrometry (AP-MS) workflow(Almasy et al., 2021b; Davies et al., 2020), in which FLAG-tagged nsp homolog constructs were transiently transfected into HEK293T cells and cells harvested 40 hours post-transfection (**Fig. 1B**). Cells were lysed and bait proteins immunopurified for FLAG-tag along with accompanying host interactors (**Fig. S1, S2**) before being reduced, alkylated, and trypsin/LysC digested. Peptides were subsequently labeled with TMTpro tags and analyzed by tandem mass spectrometry (LC-MS/MS), enabling both protein identification and relative quantification (**Tables S1, S2, Fig. S1, S2**)

**Figure 1.**
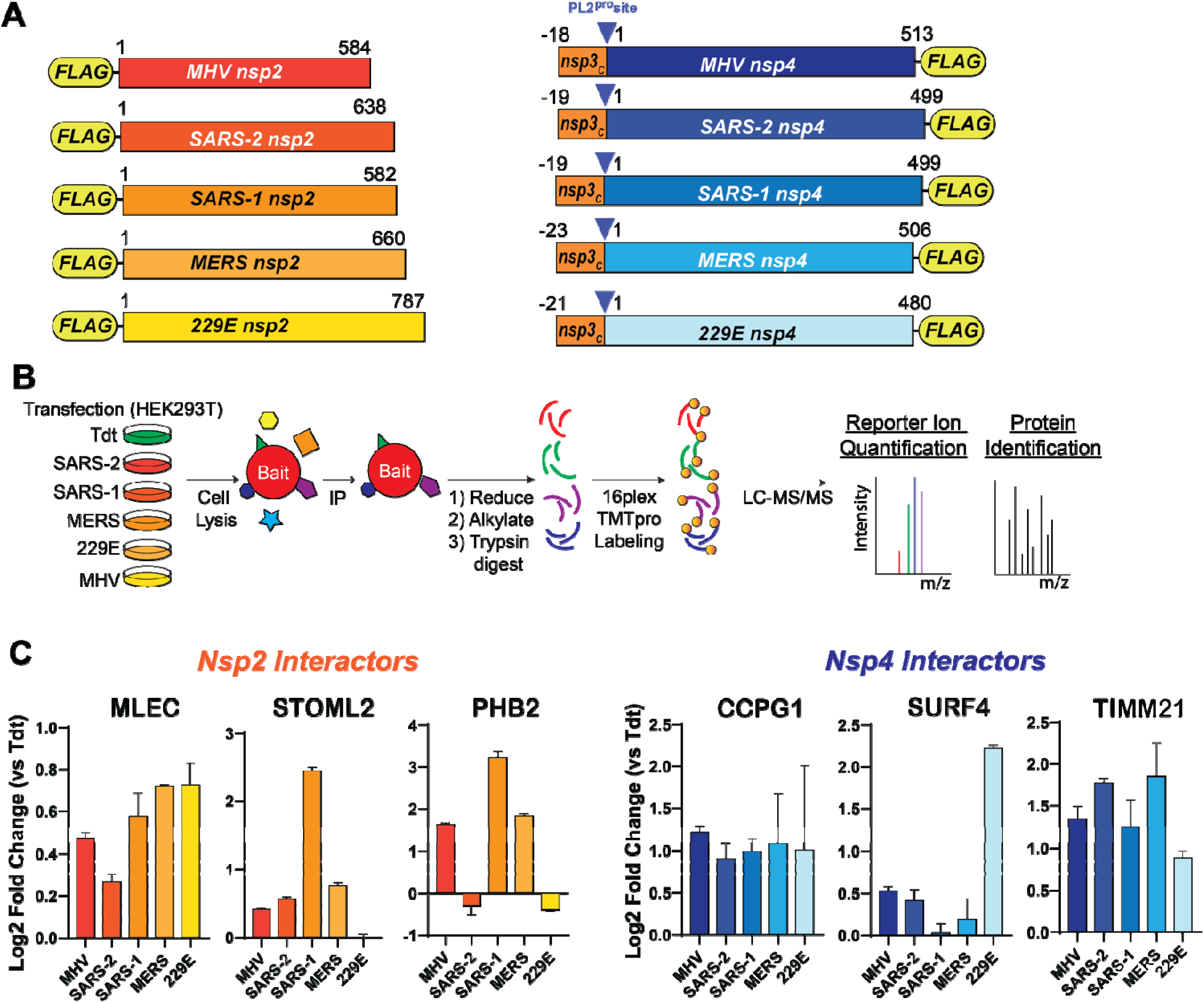
Nsp2 and nsp4 interactors are conserved across CoV homologs, including ER proteostasis factors. A. Schematic of nsp2 & nsp4 FLAG-tagged construct. The placement of the N-terminal FLAG-tag for nsp2 and C-terminal FLAG-tag for nsp4 was based on previous construct designs(Davies et al., 2020). A corresponding 18 - 23 amino acid long leader sequence from the C-terminus of nsp3, including the PL2^pro^ cleavage site, is added on the N-terminus of nsp4 constructs to improve protein biogenesis and insertion, as has been reported previously(Kanjanahaluethai et al., 2007; Oudshoorn et al., 2017). B. Schematic of comparative affinity-purification mass spectrometry (AP-MS) workflow to identify nsp-host interactors. C. Enrichment of select nsp2 and nsp4 interactors conserved across multiple CoV homologs versus negative control (Tdtomato) background. nsp2: n = 2 IP replicates/homolog in 1 MS run; nsp4: n = 2-4 IP replicates/homolog in 1 MS run. See **Figure S1, S2, Tables S1, S2** for full dataset.

We cross-referenced our previous comparative interactomics dataset with this dataset and identified several proteins of interest that were conserved across most, if not all, homologs analyzed (**Fig. S1, S2**). Several of these conserved interactors are ER proteostasis factors, such as MLEC (nsp2-interactor), CCPG1, and SURF4 (nsp4-interactors), and others are mitochondria-associated proteins, including STOML2, PHB2 (nsp2-interactors) and TIMM21 (nsp4-interactor) (**Fig. 1C**). These interactors are conserved with MHV homologs to various degrees, enabling us to utilize MHV as an accessible model BSL-2 virus to evaluate the relevance of interactors for active CoV infection.

### Conserved nsp-proteostasis interactors are pro-viral factors, including malectin

We next tested the functional relevance of conserved nsp-interactors and determine their pro- or anti-viral role in active CoV infection. We turned to the BSL-2 model betacoronavirus MHV, which has been used extensively to study CoV biology(Gosert et al., 2002; Prentice et al., 2004a; Sims et al., 2000), to screen interactors in two phases. In the first phase, priority interactors (10 nsp2 interactors, 16 nsp4 interactors) were knocked down (KD) using pooled siRNA in Delayed Brain Tumor (DBT) cells, a highly permissive cell culture model for MHV, and then infected with a previously characterized replicase reporter virus containing a genome-encoded Firefly Luciferase protein fused to the N-terminus of nsp2 (MHV-FFL2)(Freeman et al., 2014) (**Fig. 2A**). Interactors were prioritized based on conservation across multiple CoV homologs or categorization in distinct biological clusters of interest. Cells were infected at multiplicity of infection (MOI) of 0.1 and 1 plaque forming units (PFU) per cell to account for MOI-dependent effects. Scramble siRNA was used as a negative control to represent basal infection levels and Ceacam1 siRNA was used as positive control to deplete the cellular receptor for MHV. At 8 hours post infection (hpi) or 10 hpi (for MOI 1 and 0.1 respectively), cells were harvested and luminescence measured as a correlate of viral replicase protein production (**Fig. S3**).

**Figure 2.**
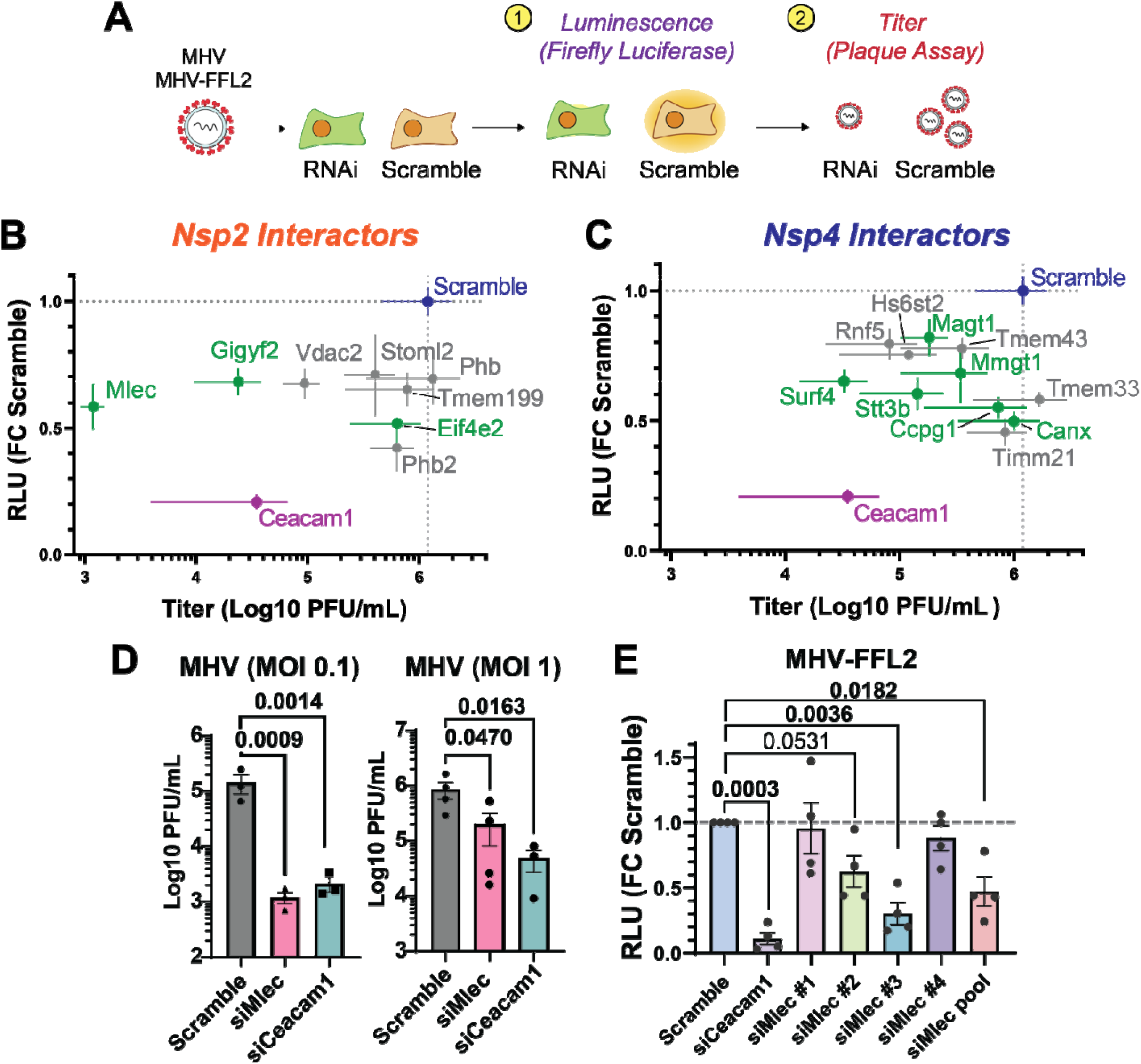
Conserved nonstructural protein (nsp) host interactors are pro-viral factors for CoV infection, including malectin (MLEC). A. Schematic of RNAi screening of prioritized interactors using a replicase reporter virus (MHV-FFL2, luminescence) or plaque assay (MHV, titer). B. Correlation plot of nsp2 interactor knockdown effect against replicase reporter virus (MHV-FFL2, luminescence) versus infectious titer (MHV, plaque). Scramble siRNA (blue) was used as basal control for comparison. siCeacam1 (MHV cellular receptor) was used as positive control (purple). Proteostasis factors are highlighted in green. Infections were performed at MOI 0.1 for 10 hpi, n=3-5. Dot represents mean, crosshatch indicates SEM. See **Figures S3, S4, S5**. C. Same as in (B), but of nsp4 interactor knockdowns. See **Figures S3, S4, S5**. D. MHV titers at MOI 0.1 or 1 in DBT cells treated with Scramble, siCeacam1, or siMlec siRNA (10 hpi), as measured by plaque assay. Paired Student’s t-test for significance, p<0.05 considered significant, n = 3 (MOI 0.1), n = 3-4 (MOI 1). E. Viral replicase reporter levels (luminescence) in DBT cells treated with Scramble, siCeacam1, individual siMlec (#1-4), or pooled siMlec siRNA, then infected with MHV-FFL2 (MOI 1, 9 hpi). Student’s t-test used for significance, with p<0.05 considered significant. n = 4. See **Figure S7**.

We identified several proteins with significant effects on the viral replicase reporter. These proteins were then advanced to the second phase of screening in which the effect of interactor KD was assessed on MHV-WT viral titers using the lower throughput plaque assay (**Fig. 2A**). The MOI and time point at which the significant effect in the luminescence reporter assay was observed were retained for the plaque assay. We identified several interactor KDs with significant reductions in titers (**Fig. S4**). To better visualize the effect of interactor KDs on both viral replicase reporter and titers, we graphed these results together as a correlation plot for MOI 0.1 (**Fig. 2B, C**) and MOI 1 (**Fig. S5**). As expected, CEACAM1 KD leads to a large reduction in both replicase reporter and titer levels. We find that almost all interactor KDs lead to a reduction in both replicase reporter and titer levels, indicating that these proteins are pro-viral factors. At MOI 0.1, there are several ER proteostasis factors which lead to substantial reductions in replicase reporter levels but with a range of effects on viral titer. For nsp2 interactors, these include the glycoprotein quality control factor malectin (MLEC), and components of the translation repression complex EIF4E2 and GIGYF2. Among nsp4 interactors were several pro-viral factors that have previously been shown to support infection of other viral families. These include MMGT1 (EMC5, a component of the ER membrane complex) (Barrows et al., 2019; Lin et al., 2019; Ngo et al., 2019), SURF4 (ER secretory cargo receptor)(Emmer et al., 2018; Kong et al., 2020; Saegusa et al., 2022; Tang et al., 2022; Yan et al., 2022), CCPG1 (ER-phagy receptor)(Smith et al., 2018), and components of the glycoprotein biogenesis factors (STT3B, MAGT1, CANX)(Marceau et al., 2016; Puschnik et al., 2017). We also tested the effects of interactor KDs on cell viability and found that none led to more than 17% reduction in viability, indicating that cell toxicity was not driving observed phenotypes (**Fig. S6**).

We found that MLEC KD yielded the largest decrease in viral titer, even surpassing the effect of CEACAM1 receptor KD. For this reason, we chose to focus on MLEC and further characterize this interaction. We verified that MLEC KD leads to titer reductions at both MOI 0.1 and 1 (10 hpi), albeit to a lesser extent with the latter, indicating that this is not a fully MOI-dependent phenotype (**Fig. 2D**). An MOI of 1 was used in all subsequent experiments unless otherwise specified.

We further validated if the reduction in MHV infection was MLEC-specific by testing the constituent individual siRNAs of the pooled siMlec treatment used previously. We find that at least two of the four siMlec (#2, 3) lead to reductions in viral replicase reporter levels (as measured with MHV-FFL2) and that endogenous MLEC protein levels are reduced by ∼80% compared to Scramble (**Fig. 2E, Fig. S7**). Together, these results support a MLEC-specific phenotype and show that MHV is highly dependent on MLEC as a pro-viral factor for infection.

### Nsp2 interacts with MLEC-associated protein complexes during infection

MLEC is an ER-resident glycoprotein quality control factor that binds with high affinity to diglucosylated moieties on protein-linked oligosaccharides in the ER lumen (Schallus et al., 2010, 2008). These oligosaccharides are typically rapidly trimmed from triglucosyl to monoglucosyl moieties by Glucosidase I and II, but diglucosylated proteins may result from mistrimming events likely related to misfolding(Tannous et al., 2015). MLEC expression is upregulated during ER stress, when misfolded proteins accumulate, and has been found to retain diglucosylated glycoproteins in the ER lumen to prevent the secretion of misfolded glycoprotein(Chen et al., 2011; Galli et al., 2011). MLEC has also been shown to associate with ribophorin I (RPN1)(Qin et al., 2012; Ramírez et al., 2019; Takeda et al., 2014), a component of the OST complex, indicating a tight and early linkage between the initial addition of the triglucosylated oligosaccharide by the OST complex and the surveillance function of MLEC (**Fig. 3A**).

**Figure 3.**
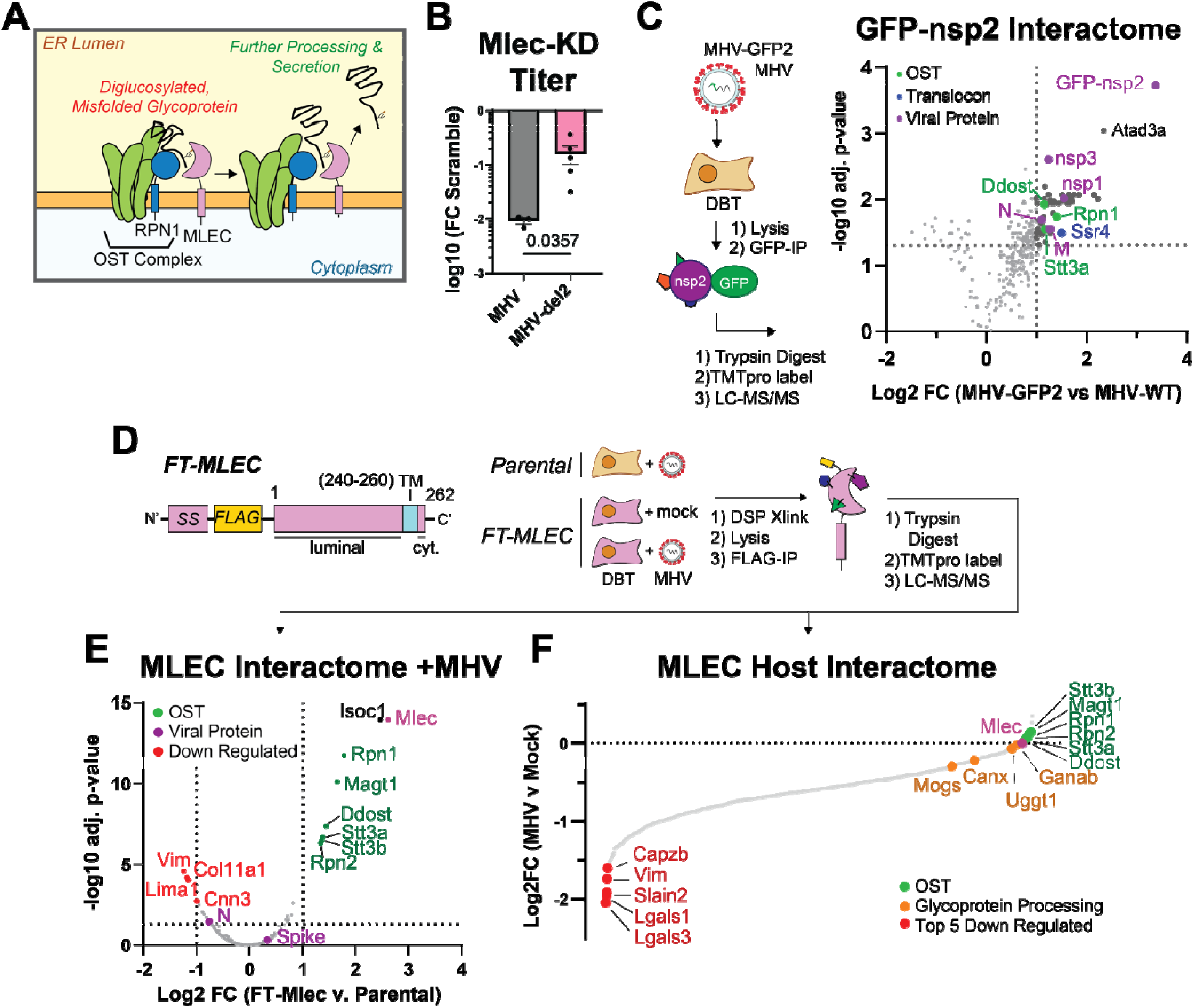
Nsp2 maintains a genetic interaction with MLEC and protein-protein interactions with MLECassociated protein complexes during infection. A. Schematic of the role of MLEC in glycoprotein quality control. B. Comparison of MHV-WT vs. MHV-del2 (nsp2 deleted) infectious titers in siMlec vs. Scramble-treated DBT cells, expressed as Log10 fold change. MHV-WT (MOI 0.1, 10 hpi) or MHV-del2 (MOI 0.5, 10 hpi). n = 3-5, with Mann-Whitney T-test for significance, p <0.05 considered significant. See **Figure S8**. C. Volcano plot of MHV GFP-nsp2 interactome during infection (MOI 1, 9 hpi). Comparison to MHV-WT i fected cells was used as a background control for the GFP-IP. Linear filters of Log2 fold change > 1 and adjusted p-value > 0.05 were applied. Viral proteins (purple), OST complex factors (green), and translocon factors (blue) are annotated. MHV-GFP2 (n = 8 IPs), MHV-WT (n = 7 IPs) combined in 1 MS run. One-way ANOVA with multiple testing corrections was used for significance. See **Figure S9, Table S3**. D. Schematic of FT-MLEC construct design and IP-MS experimental design. FT-MLEC construct is composed of an N-terminal signal sequence (SS), followed by a FLAG-tag, and the MLEC luminal domain. There is a 10-residue-long transmembrane helix with a short 2 amino acid long cytosolic C-terminal tail. Stably expressing FT-MLEC or WT DBT cells were infected with MHV (MOI 2.5, 9 hpi) and subjected to DSP (0.5 mM) crosslinking, before lysis and FLAG IP. Elutions were reduced, alkylated, trypsin digested, and analyzed by LC-MS/MS. Parental DBT and MHV (n = 4 IPs), FT-MLEC DBT and mock (n = 6 IPs), FT-MLEC DBT and MHV (n = 6 IPs/condition), pooled into 1 MS run. See **Figure S10**. E. Volcano plot of FT-MLEC interactors during MHV infection. Samples were normalized to total peptide amounts. Linear filters of Log2 fold change > 1 or < -1 and adjusted p-value > 0.05 were applied. Viral proteins and MLEC (purple), downregulated (red), and OST complex factors (green) are annotated. T-test with multiple testing corrections for significance. See **Figure S11, Table S4**. F. Waterfall plot of FT-MLEC interactome comparing MHV versus mock infection. Samples were normalized to FT-MLEC bait levels. Components of the OST complex (green), the glycoprotein processing pathway (orange), and the five most downregulated interactors (red) are annotated. See **Figure S11**.

As it was surprising that nsp2, a non-glycosylated, cytoplasmic protein, would interact with MLEC, an integral ER membrane protein with a short two amino acid cytoplasmic tail(Galli et al., 2011; Ramírez et al., 2019), we assessed a broader genetic interaction between nsp2 and MLEC. To this end, we compared the effect of MLEC-KD on WT MHV infection versus infection with an MHV strain lacking nsp2 (MHV-del2)(Graham et al., 2005) (**Fig. 3B, Fig. S8**). MLEC KD leads to only an order of magnitude (log_10_ PFU/mL) reduction in titer of MHV-del2, as opposed to the two orders of magnitude reduction of MHV-WT, indicating that nsp2 and MLEC cooperate to promote viral infection.

We next determined if nsp2 forms a PPI with MLEC during active infection. We profiled the nsp2 interactome by infecting DBT cells (MOI 1) with an MHV strain genetically encoding GFP fused to the N-terminus of nsp2 (MHV-GFP2), which has been previously characterized(Freeman et al., 2014). Infection with WT MHV was used as a background control for the GFP IP. At 9 hpi, cells were harvested and lysed with gentle lysis buffer to maintain native interactions, then immunoprecipitated for GFP-nsp2. Elutions were reduced, alkylated, trypsin digested, and TMTpro labeled prior to LC-MS/MS analysis as described previously (**Fig. 3C, Table S3, Fig. S9**).

We found that nsp2 interacts with several viral proteins, including the structural nucleocapsid (N) and membrane (M) proteins, and the nonstructural proteins nsp1 and nsp3. We also observe several mitochondrial interactors, most notably ATAD3A, which acts as a scaffold protein on the surface of the mitochondria to tether the outer and inner mitochondrial membranes together(Arguello et al., 2021; Zhao et al., 2019). This interaction may suggest a role for nsp2 in modifying mitochondrial activity, as previously suggested(Davies et al., 2020).

Excitingly, we identify several interactions between nsp2 and the OST complex (RPN1, STT3A, DDOST) and SSR4 (a component of the translocon complex). While MLEC itself was not identified, these interactors are previously known to associate with MLEC(Qin et al., 2012; Ramírez et al., 2019; Takeda et al., 2014). These proteins themselves are integral ER membrane proteins with minimal cytosolic domain presence, likely indicating there may be some intermediary protein facilitating these interactions. Nsp2 has been shown to localize to RTC sites from the cytoplasmic face(Freeman et al., 2014; Hagemeijer et al., 2009). By extension, these interactions between the OST complex and MLEC may be concentrated at these replication sites, pointing towards a broader link between nsp2 and glycoprotein processing machinery during infection.

We next asked whether MLEC interacts with nsp2 or any other viral proteins during CoV infection, and if MHV infection alters the interactome of MLEC. To this end, we generated DBT cells stably expressing an N-terminal FLAG-tagged construct of MLEC (FT-MLEC) (**Fig. 3D**). We validated that the signal sequence of this construct is properly cleaved (as evident by the immunoblotting reactivity to M1 FLAG antibody) and that FT-MLEC localizes to the ER, as shown by colocalization IF with PDIA4, an ER marker (**Fig. S10**). Viral titers in FT-MLEC DBT cells are similar to WT DBT cells, confirming that overexpression of FT-MLEC in these cells does not alter MHV infectivity (**Fig. S10**).

To characterize the interactome of MLEC during infection, we infected FT-MLEC DBT cells with MHV (MOI 2.5) for 10 hpi (**Fig. 3D**). Parental DBT cells were infected to serve as a background control for FLAG-IP. Mock infected FT-MLEC DBT cells served as a basal control to differentiate how MHV infection alters the MLEC interactome. Prior to harvest, infected cells were subjected to crosslinking with dithiobis(succinimidyl propionate) (DSP, 0.5 mM) to capture potentially transient interactions characteristic of lectin-client interactions and then lysed and immunoprecipitated for FLAG-tag. Elutions were subsequently trypsin digested, labeled with TMTpro, pooled, and analyzed by tandem mass spectrometry (LC-MS/MS) as described previously.

When comparing FT-MLEC versus Parental DBT cells to identify MLEC-interacting proteins during infection, several members of the OST complex (RPN1, RPN2, MAGT1, DDOST, STT3A, STT3B) were significantly enriched as expected, given they are canonical MLEC interactors (**Fig. 3E, Table S4, Fig. S11**). The MHV Spike protein was positively but only modestly enriched.

We also examined the effect of MHV infection on the host interactome of MLEC (**Fig. 3F**). Comparing FT-MLEC IPs from cells either mock or MHV infected (normalized to FT-MLEC levels, **Fig. S11**) revealed a large loss of the basal MLEC interactome during infection. Some of the most downregulated proteins include CAPZB, VIM, SLAIN2, and LGALS1/3. LGALS proteins (galectins) are a group of lectins that bind beta-galactosides and typically reside in the cytosol, though they can be secreted through transport into the ER-Golgi intermediate compartment (ERGIC)(Zhang et al., 2020). There is also a more modest downregulation of interactions with glycoprotein processing factors, including MOGS (glucosidase I, GluI) and to a lesser extent GANAB (glucosidase II, GluII). GluI acts upstream of MLEC to trim the triglucosyl oligosaccharide on glycoproteins to a diglucosyl moiety, whereas GluII acts downstream to trim the diglucosyl oligosaccharide to a monoglucosyl group(D’Alessio et al., 2010; Deprez et al., 2005; Totani et al., 2006). From there, client glycoproteins enter the calnexin (CANX) folding cycle, which is partially regulated by UGGT1(Adams et al., 2020; Ellgaard and Helenius, 2003; Guay et al., 2023; Hammond et al., 1994). Both CANX and UGGT1 are modestly downregulated interactors during MHV infection, suggesting a possible decoupling of early and late glycoprotein quality control systems.

In contrast, OST complex interactors (RPN1/2, STT3A/B, MAGT1, DDOST) are maintained and even slightly upregulated during infection. These results indicate that MHV infection induces a reorganization of the MLEC host interactome in which some canonical interactions are retained (the OST complex), while other components of the glycoprotein biogenesis pathway and other lectin family proteins dissociate.

In summary, nsp2 coordinates with MLEC to exert its pro-viral function and interacts with MLEC-associated proteins during infection. Furthermore, MHV infection seems to drastically alter the interactome of MLEC, stabilizing OST interactions while weakening other lectin interactors.

### MLEC promotes early CoV replication events and viral protein biogenesis

We next sought to understand at which stage in the viral replication cycle MLEC promotes infection, (**Fig. 4A**). To test if MLEC promotes viral entry, we measured abundance of viral positive-sense genomic RNA, *(+)gRNA*, present within the cell early in MHV infection (MOI 1, 2 hpi) via quantitative reverse transcription PCR (RT-qPCR) (**Fig. 4B**, **Table 1**). KD of the MHV cellular receptor CEACAM1 leads to a reduction in *(+)gRNA* levels compared to Scramble control, whereas siMlec actually leads to an increase in intracellular *(+)gRNA*. This finding indicates MLEC does not promote viral entry and genome release.

**Figure 4.**
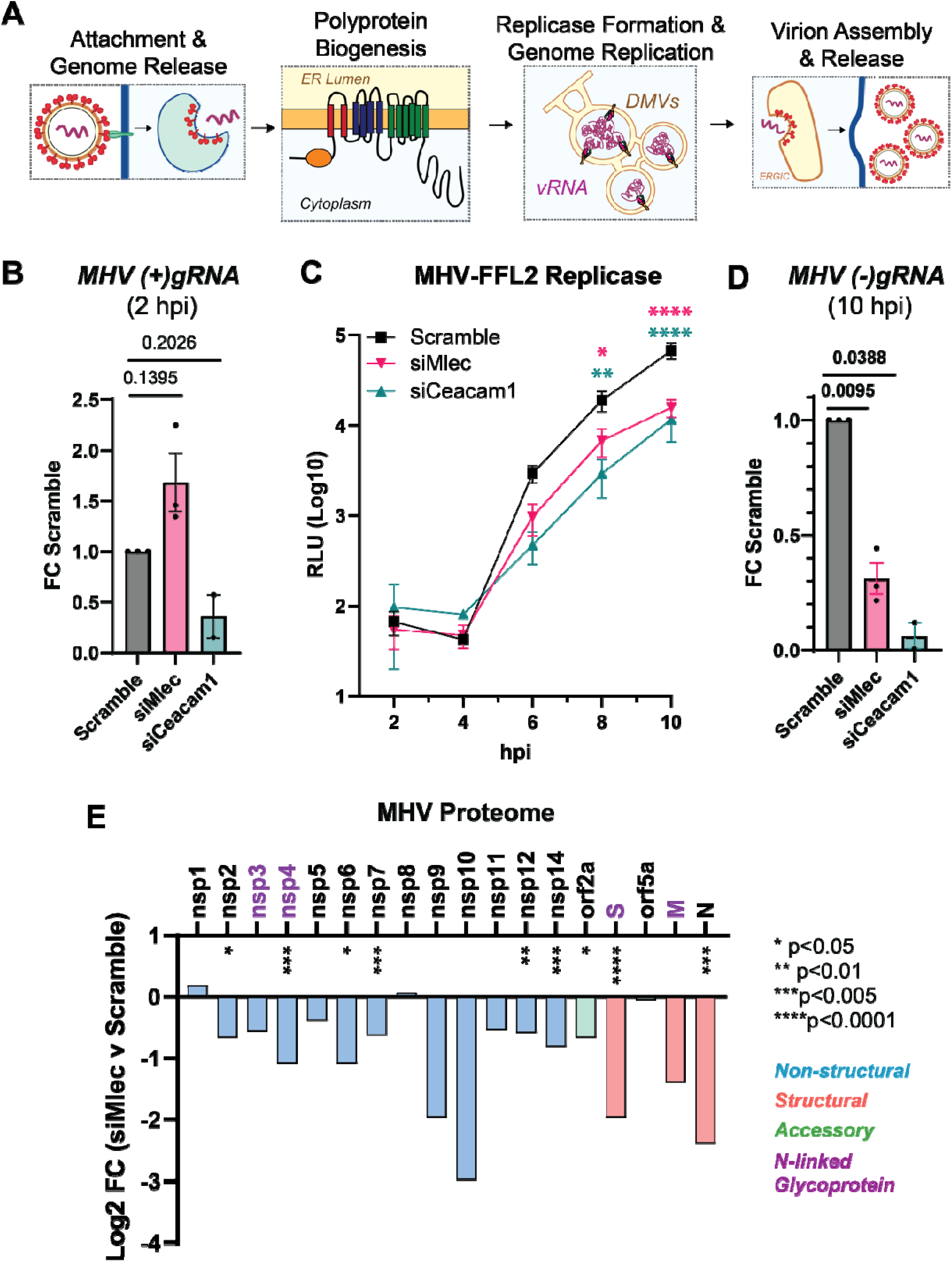
MLEC promotes early CoV replication events and viral protein biogenesis. A. Schematic of main events during the CoV replication cycle. Viral RNA (vRNA), double membrane vesicle (DMV), ER-Golgi Intermediate Compartment (ERGIC). B. Abundance of positive-sense MHV genomic RNA (*(+)gRNA*) as measured by RT-qPCR (MHV WT, MOI 1, 2 hpi) and normalized to *Rpl13a* transcript levels. Paired Student’s T-test for significance, p<0.05 considered significant, n = 2-3. C. MHV-FFL2 replicase reporter time course during infection from 2 – 10 hpi (MOI 1). Mock infected cells used as background luminescence control. Two-way ANOVA, Benjamini, Krieger, Yekutieli multiple comparisons, n = 3, * p<0.05, ** p<0.01, ***p<0.005, ****p<0.0001, color coding represents results of siMlec (pink) or siCeacam1 (teal) compared to Scramble control (black). D. Abundance of negative-sense MHV genomic RNA (*(-)gRNA*) as measured by RT-qPCR (MHV WT, MOI 1, 10 hpi). Paired Student’s T-test, p<0.05 considered significant, n = 2-3. See **Figure S12**. E. Relative abundance of intracellular viral proteins during MHV infection in DBT cells treated with Scramble or siMlec siRNA (MOI 1, 10 hpi), measured by quantitative proteomics. Purple protein names denote viral glycoproteins. Spike protein (S), membrane protein (M), nucleocapsid protein (N). One-way ANOVA with multiple testing correction. * p<0.05, ** p<0.01, ***p<0.005, ****p<0.0001, n = 6-7 biological replicates/condition across 2 MS runs. See **Figure S13, Table S5**.

**Table 1.**
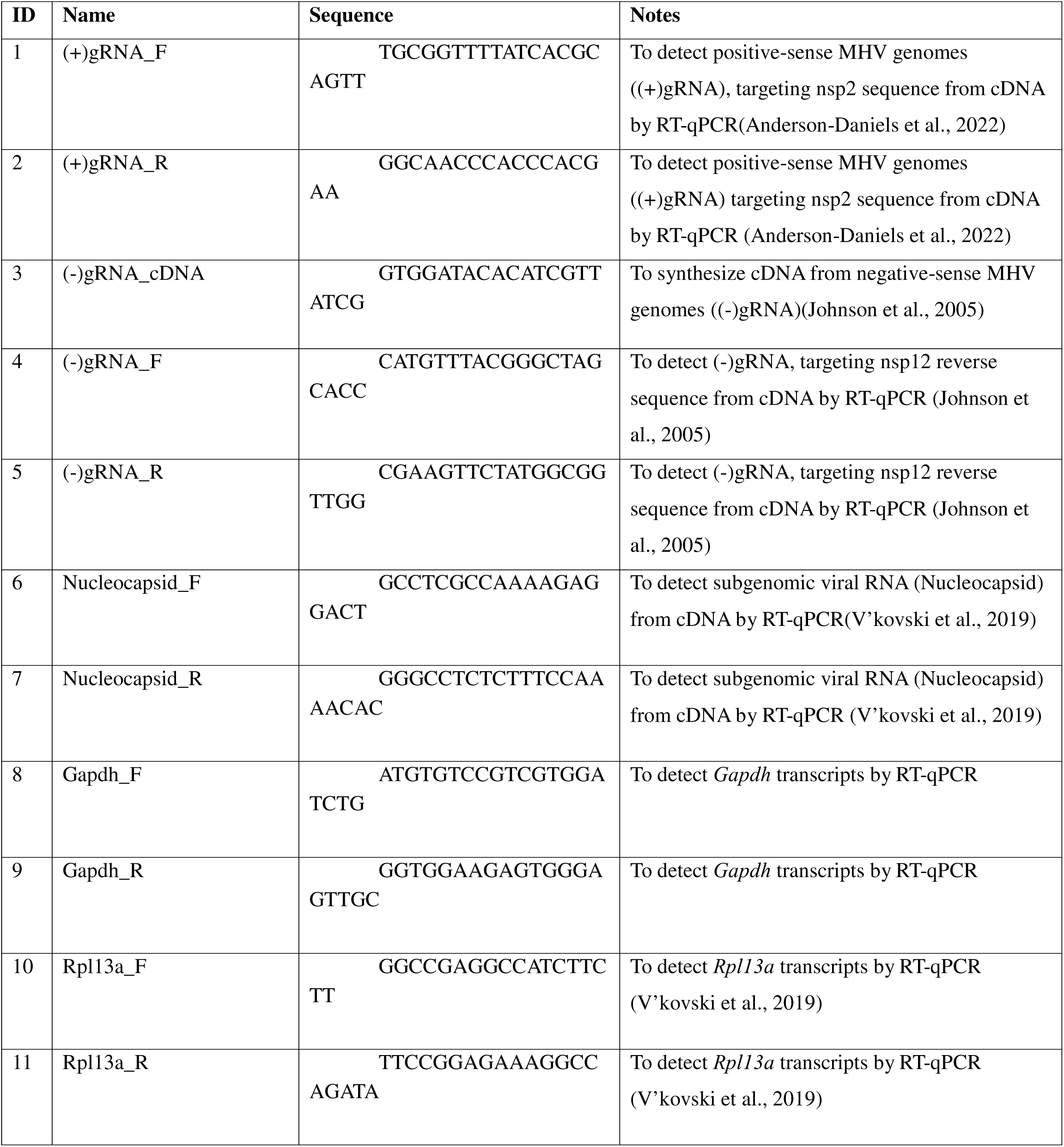
Primers used in this study.

Next, we tested whether MLEC promotes viral polyprotein production using the MHV-FFL2 virus, which reports on the abundance of nsp2 as a correlate of replicase protein production. DBT cells were treated with scramble, siCeacam1, or siMlec and then infected with MHV-FFL2 (MOI 1) and lysates harvested every two hours, from 2 to 10 hpi (**Fig. 4C**). Early during infection, at 2 and 4 hpi, luminescence levels are low and relatively equal amongst conditions. At 6 hpi, siMlec and siCeacam1 treatment stunted luminescence production compared to the scramble control, and this difference continued to widen significantly at 8 and 10 hp. These results indicate MLEC promotes early replicase protein production.

We hypothesized then that a reduction in replicase proteins would in turn result in a loss of viral genomic RNA production. To test this outcome, we used RT-qPCR to measure the abundance of negative-sense viral gRNA, *(-)gRNA*, which is the biochemical intermediate of (+)gRNA replication, at 10 hpi (**Fig. 4D**, **Table 1**). As expected, MLEC-KD significantly reduces *(-)gRNA* levels compared to Scramble control. We also find that *(+)gRNA* and *Nucleocapsid* transcripts are significantly reduced at 10 hpi, corroborating this result (**Fig. S12**).

Since MLEC KD reduces nsp2 levels, as seen in the MHV-FFL2 time course, we turned to quantitative proteomics to assess how MLEC KD affects the abundance of the rest of the MHV proteome. Comparing the siMlec vs. Scramble conditions revealed a global reduction in the total viral proteome, with significant decreases in structural, non-structural, and accessory proteins (**Fig. 4E, Table S5, Fig. S13**). There are significant reductions in several viral glycoproteins, including Spike, nsp4, and nsp6. However, MLEC KD also affects non-glycosylated protein levels, such as nucleocapsid and nsp2. Considering that nsps are translated during infection and that structural proteins are translated from subgenomic RNAs (sgRNAs) produced during viral genomic replication, the reduction in nsp levels with MLEC KD likely spurs the subsequent drop in structural protein abundance. As such, we conclude that MLEC promotes nsp protein production and MLEC KD leads to reduced nsp abundance, which cascades into loss of viral genome replication and subsequent defect in viral titers.

### MLEC mediates viral replication through the glycoprotein biogenesis pathway

Since MLEC acts as a quality control factor in the *N*-linked glycoprotein biogenesis pathway, we wanted to test whether MLEC promotes viral protein production through this function. We used the small molecule NGI-1, which inhibits the glycan transfer activity of the OST complex, upstream of GluI(Lopez-Sambrooks et al., 2016) (**Fig. 5A**). If MLEC influenced MHV replication through a mechanism independent of the glycoprotein biogenesis, then the inhibitor and MLEC KD should both suppress viral replication, producing an additive effect. DBT cells were transfected with Scramble siRNA (negative control), siCeacam1 (positive control), or siMlec and then treated with 5 µM NGI-1 or DMSO at the start of infection with MHV-FFL2 (MOI 1, 10 hpi). This dosage and timing were chosen to partially inhibit the OST complex without fully ablating viral infection, as NGI-1 has been shown previously to be a potent positive-sense RNA virus inhibitor(Puschnik et al., 2017) (**Fig. S14**). Lysates of infected cells were analyzed by Western blot to verify MLEC KD and probe for ERDJ3, an ER-resident glycosylated chaperone, to confirm *N*-linked glycosylation inhibition (**Fig. 5B, Fig. S14**). We found that the reduction in viral replicase reporter levels upon NGI-1 treatment were comparable between Scramble and MLEC KD cells, indicating that there is not an additive effect of upstream OST inhibition and MLEC KD. This result confirms that MLEC promotes viral replication through its canonical N-glycan-dependent function (**Fig. 5C**). In contrast, CEACAM1 KD combined with NGI-1 treatment leads to a slightly increased suppression of MHV-FFL2 replicase reporter levels compared to the scramble siRNA control, albeit this change is highly variable and does not reach significance.

**Figure 5.**
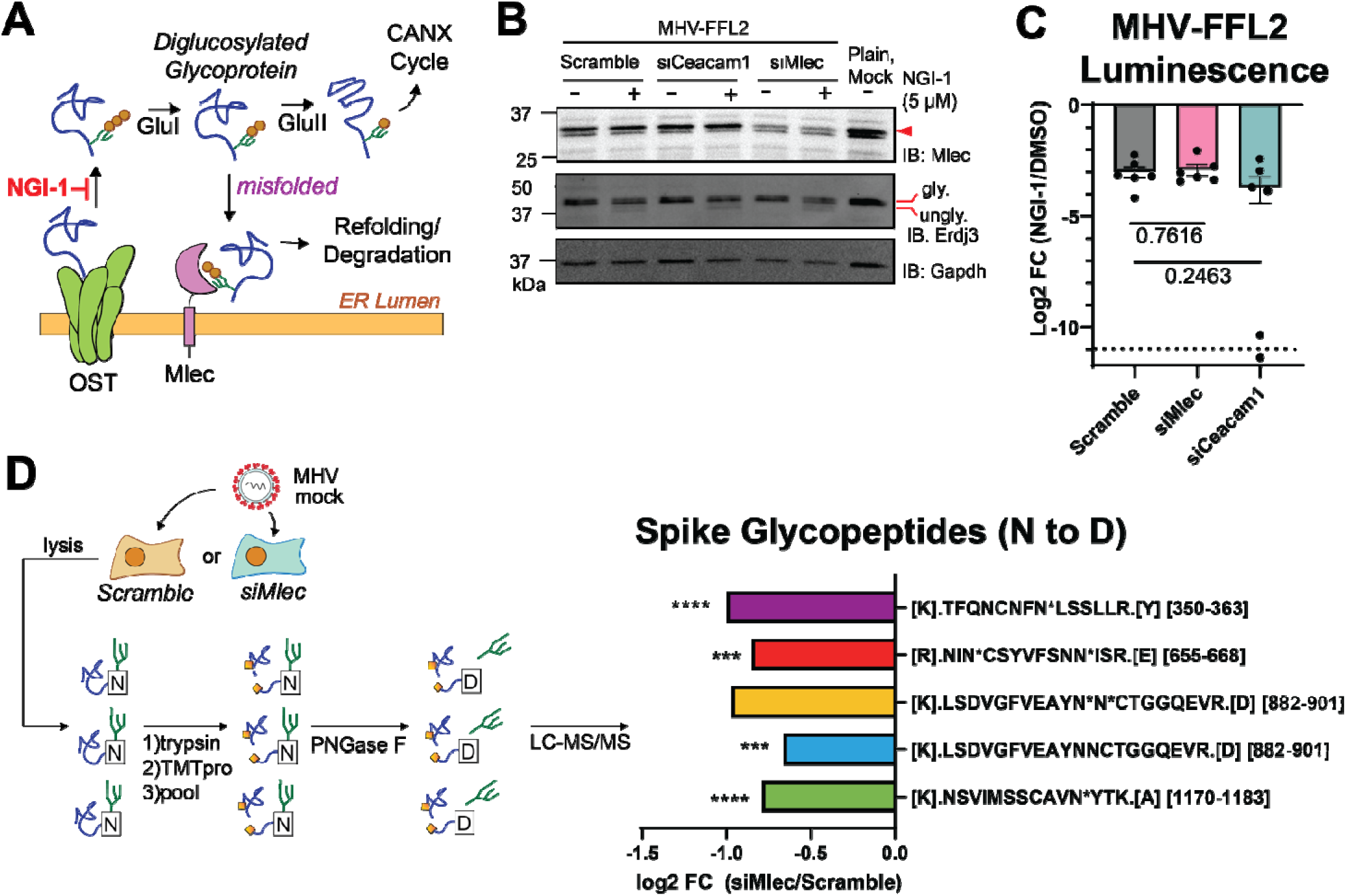
MLEC mediates replication through the glycoprotein quality control pathway. A. Schematic of glycoprotein biogenesis and quality control pathway. NGI-1 is an OST inhibitor that blocks *N*-linked glycosylation(Lopez-Sambrooks et al., 2016). B. Representative Western blot of DBT lysates after OST inhibition and MLEC KD during MHV-FFL2 infection (MOI 1, 10 hpi). Blotting for glycosylated and non-glycosylated ERDJ3 served as a marker of *N*-linked glycosylation inhibition. Lysates were matched with the corresponding luminescence reading. See **Figure S14**. C. Levels of MHV-FFL2 replicase reporter with NGI-1 or DMSO treatment at the time of infection in cells with Scramble, CEACAM1, or MLEC KD in the same lysates analyzed in (B). One replicate of siCeacam1 fell below the limit of detection (dotted line). Mock-infected cells were used as a control for background luminescence. Student’s T-test for significance, with p<0.05 considered significant, n = 6. See **Figure S14**. D. Abundance of *N*-linked glycosylated Spike peptides in siMlec versus Scramble-treated DBT cells. Cells were treated with siRNA and infected with mock or MHV (MOI 1, 10 hpi). Lysates were trypsinized, TMTpro labeled, pooled, then treated with PNGase F to cleave oligosaccharides, resulting in a deamidation mass shift (N to D) detectable by LC-MS/MS. Peptide sequence is listed, with “N*” denotating a deamidation modified Asn in a canonical glycosylation sequon (N-X-S/T, where X is not Proline). Peptides where the modified Asn could not be resolved lack the “N*” annotation. Peptide position within the protein is in brackets. One-way ANOVA, adjusted p-value. * p<0.05, ** p<0.01, ***p<0.005, ****p<0.0001, n = 4/condition, 1 MS run. See **Figure S15, Table S6**.

We then asked how the knockdown of MLEC might affect the *N*-linked glycosylation status of viral proteins during infection, as MLEC KD may lead to disruptions in the glycoprotein biogenesis pathway and decreased glycosylation of viral glycoproteins. To assess the glycosylation status of viral proteins, we treated DBT cells with Scramble or siMlec siRNA and then infected cells with MHV (MOI 1, 10 hpi). Lysates were trypsin digested, TMTpro labeled, and pooled as before. Labeled peptides were then treated with PNGase F to trim off *N*-linked oligosaccharides, leading to an Asn to Asp conversion and a mass shift (deamidation, +0.984 Da) detectable by LC-MS/MS analysis(Adams et al., 2020; Guay et al., 2023) (**Fig. 5D, Table S6, Fig. S15**). Peptides were then filtered for deamidation modifications and verified to contain canonical *N*-linked glycosylation site sequons (N-X-S/T, where X is not Proline)(termed glycopeptides).

We were unable to detect glycopeptides from nsps but identified five glycopeptides from the Spike protein (**Fig. 5D, Fig. S15B**). All five showed a decrease in abundance in siMlec versus scramble siRNA treated cells, with four being statistically significant. While we cannot ascertain the effect on nsp glycopeptides with this data, these results further support a role for MLEC promoting viral protein biogenesis through the glycoprotein quality control pathway.

This mass spectrometry data set also revealed general alterations to the cellular proteome with MLEC KD and MHV infection. In uninfected cells, MLEC KD leads to relatively little proteome-wide changes, with MLEC being the only protein significantly downregulated and no other proteins significantly upregulated, supporting the specificity of MLEC KD in MHV suppression (**Fig. S15C**). To determine whether MLEC KD alters general host proteostasis, we further examined the levels of protein markers of stress pathways based on previous gene pathway definitions(Davies et al., 2023; Grandjean et al., 2019; Shoulders et al., 2013) (**Fig. S15D**). We find that there are modest but significant increases in protein levels associated with the Heat Shock Factor 1 (HSF1) pathway, while the Unfolded Protein Response (UPR) pathways are largely unmodified.

We also probed the effect of MLEC KD on endogenous protein glycosylation. We find that there is only a small increase in abundance of glycopeptides, including those associated with the ribosome (Rpl3, Rpl5), a cytoskeletal protein (Rdx), the integrin Itgb1, and the ER-resident chaperone Serphinh1 (**Fig. S15E-G**).

As expected, in Scramble-treated cells, MHV infection leads to large alterations in the cellular proteome (**Fig. S15H**), and MLEC KD attenuates these changes and viral protein abundance (**Fig. S15I**).

### MLEC promotes SARS-CoV-2 replication

While MHV is a longstanding beta-coronavirus model, we sought to test whether MLEC also promotes replication of the human pathogen SARS-CoV-2. In lieu of BSL-3 facilities, we turned to a previously characterized SARS-CoV-2 replicon plasmid system(Taha et al., 2023) which expresses all SARS-CoV-2 proteins with a secreted nanoLuciferase (sec-nLuc) reporter in place of Spike. HEK293T cells were transfected with siMLEC siRNA and either the Delta or Omicron SARS-CoV-2 replicon plasmid. GFP expression plasmid and Scramble siRNA served as controls. After 72 h post-transfection (hpt), sec-nLuc luminescence activity was measured as a reporter for SARS-CoV-2 replication (**Fig. 6A**). Co-transfection with siMLEC leads to a significant decrease in Delta replication compared to Scramble. MLEC knockdown also suppresses Omicron replication, though not significantly. Consistent with previous reports(Taha et al., 2023), the Delta replicon produced higher reporter levels than Omicron. To normalize for these differences in magnitude, we calculated the fold change of siMLEC versus Scramble treatment and found that MLEC knockdown suppresses replication to 43 and 41 % of WT levels of Delta and Omicron respectively (**Fig. 6B**). MLEC protein knockdown was verified by Western blot (**Fig. S16A**).

**Figure 6.**
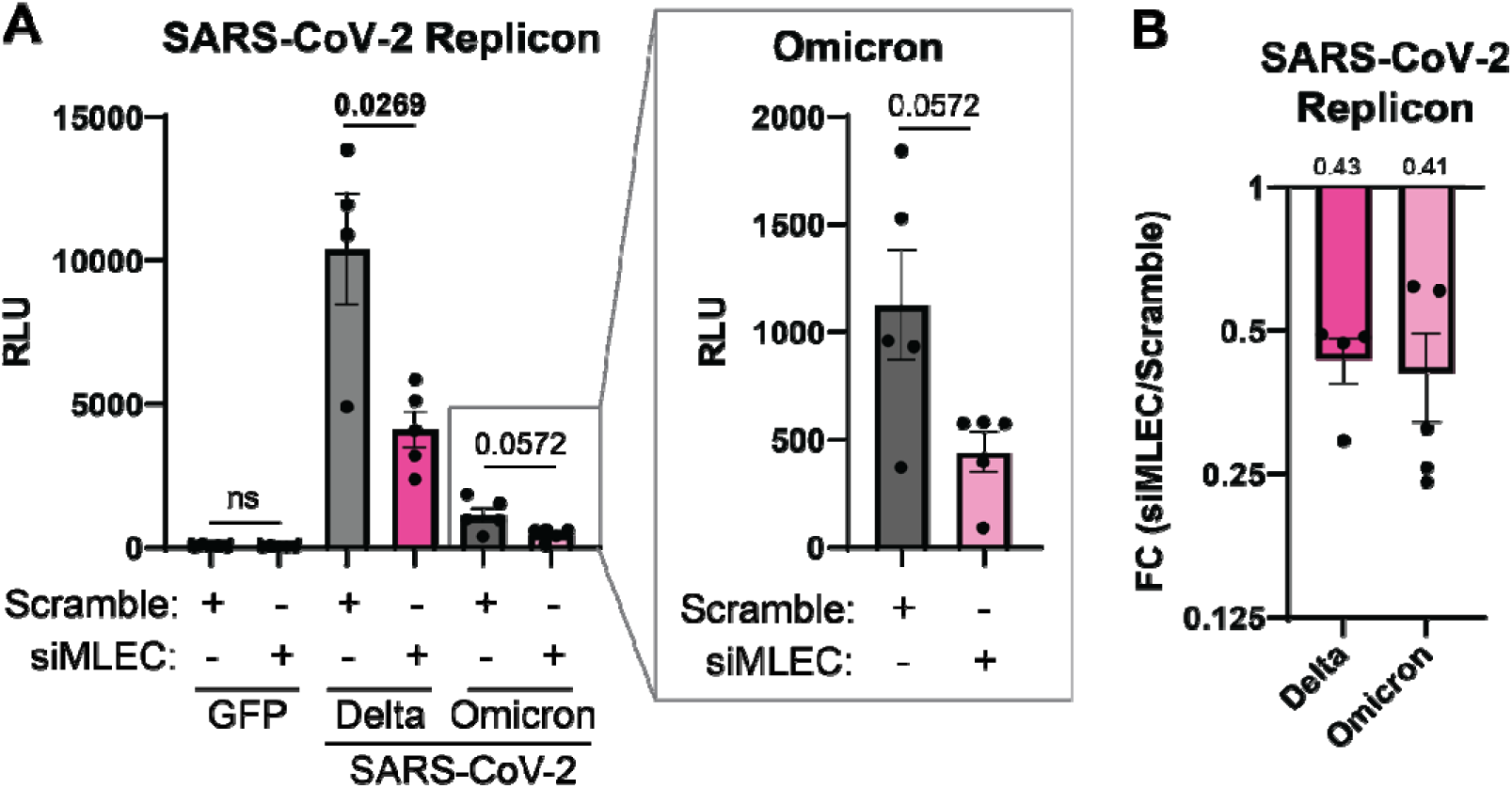
MLEC knockdown suppresses SARS-CoV-2 replication. A. Luminescence levels of a secreted SARS-CoV-2 replicon nano-Glo luciferase (sec-nLuc) reporter(Taha et al., 2023) for Delta or Omicron in HEK293T cells treated with siMLEC or Scramble. GFP was used as a negative control. Omicron data is displayed in subset box. Delta, n = 4; Omicron, n = 5; Student’s t-test for significance, with p < 0.05 considered significant. Bars represent mean values, ± SEM. See **Figure S16**. B. Data from (A) presented as a fold change comparison of siMLEC vs. Scramble treatment with respective SARS-CoV-2 replicon. Mean fold change value is annotated above the graph (0.43 and 0.41 respectively). Bars represent mean values, ± SEM.

To control for the possibility that MLEC knockdown reduces nLuc secretion independent of effects on replicon production, we also measured the abundance of internal (non-secreted) nLuc from the lysates of the same samples (**Fig. S16B-C**). The internal nLuc luminescence is similarly reduced as observed for secreted nLuc, indicating that MLEC promotes SARS-CoV-2 replication.

## Discussion

This study aims to characterize conserved protein-protein interactions (PPIs) between CoV nsps and host proteostasis factors, building on our previous work(Davies et al., 2020). By expanding the panel of nsp homologs tested, we were able to identify interactors conserved with additional CoV strains, including MERS-CoV, hCoV-229E, and MHV (**Fig. 1**). MHV was then used as a model virus to determine the pro- or anti-viral roles of conserved interactors. We identified several conserved ER proteostasis factors with pro-viral roles, most notably the glycoprotein quality control factor Malectin (MLEC), which leads to a drastic reduction in viral titers (**Fig. 2**). We also found GIGYF2-KD strongly suppressed MHV infection, despite GIGYF2 not interacting with MHV nsp2 (**Fig. S1D**), highlighting the importance of proteostasis factors in infection regardless of direct PPIs.

Interrogating the nsp2-MLEC interaction, we found that nsp2 and MLEC maintain a genetic interaction and that nsp2 interacts with MLEC-associated proteins such as the OST complex during infection. MHV infection also remodels the MLEC interactome, retaining MLEC-OST complex interactions while diminishing MLEC interactions with downstream glycoprotein processing factors (**Fig. 3**). We show that MLEC KD leads to defects in viral nsp production, which likely induces subsequent reductions in viral genome replication and structural protein production (**Fig. 4**). We find that MLEC promotes viral replication through the glycoprotein biogenesis pathway and MLEC KD leads to a decrease in viral glycopeptides during infection (**Fig. 5**). Lastly, using a replicon system, we show that MLEC promotes SARS-CoV-2 replication (**Fig. 6**). Our findings are summarized in **Figure 7**.

**Figure 7.**
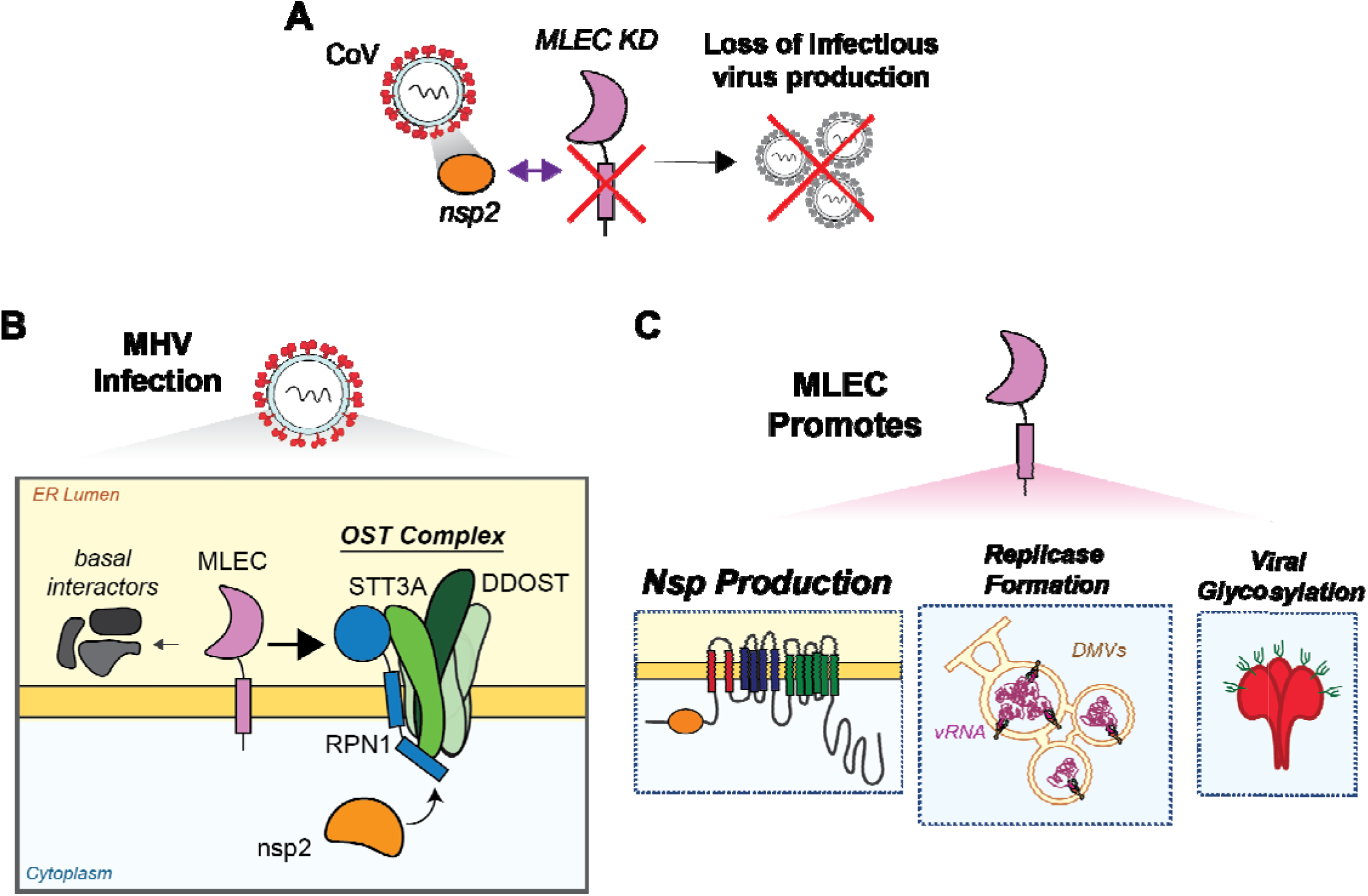
Summary of the role of MLEC in coronavirus replication. A. CoV nsp2 homologs interact with MLEC when measured by protein expression in HEK293T cells. Knockdown of MLEC results in loss of infectious MHV production and SARS-CoV-2 replication. B. During MHV infection, MLEC loses many basal interactors while maintaining interactions with the OST complex. Nsp2 interacts with core components of the OST, including RPN1, STT3A, and DDOST. C. MLEC promotes biogenesis of CoV nsps and subsequent replicase formation. In addition, MLEC knockdown results in suppression of Spike glycopeptides, likely through the canonical role of MLEC in the glycoprotein quality control pathway.

A limitation of our study is the initial overexpression of individual proteins for AP-MS, in which we find some variation between our data with other AP-MS studies. We sought to overcome these variations by focusing on conserved interactors and testing interactions in a live infection context.

There are several potential mechanisms by which MLEC mediates CoV infection. We were unable to detect a direct PPI between MLEC and nsps during MHV infection, so the pro-viral effect may not be mediated through a direction interaction. It is possible that the conditions used here were not conducive to detecting these PPIs or that interactions are more prominent in earlier timepoints of infection as opposed to the one tested (10 hpi). Further work should test alternative conditions and timepoints.

Alternatively, nsp2 and MLEC are likely connected through an indirect interaction. We find that nsp2 interacts with several OST complex members, including DDOST, STT3A, and RPN1, though whether this is as part of the uncleaved Orf1a polyprotein during co-translational ER translocation or as an individual protein is unclear. RPN1 is a bona fide interactor of MLEC, which has been shown by both IP and colocalization studies(Qin et al., 2012; Takeda et al., 2014), as well as in a cryo-EM structure which reveals that MLEC directly engages with RPN1 in the OST complex(Ramírez et al., 2019). Further, MHV infection retains the association of MLEC with the OST complex while titrating off other interactors, potentially leading to more efficient glycoprotein biogenesis. Given that nsp2 has previously been shown to localize with RTCs during infection(Freeman et al., 2014; Hagemeijer et al., 2009) and we detect nsp2 interactions with nsp3 (an RTC marker), we propose that the nsp2-OST-MLEC interaction axis leads to enhanced viral glycoprotein production around RTCs, most prominently the biogenesis of nsps.

This is in line with previous proximity-labeling work that showed host cell translational machinery locate near nsp2 and RTCs(V’kovski et al., 2019). Concurrently, knockdown of MLEC leads to impediment of nsp production and aberrant glycosylation of other viral proteins like Spike, though it should be noted that the decrease in Spike glycopeptides is compounded by the overall decrease in Spike protein. Given that MLEC is pro-viral in a SARS-CoV-2 replicon model lacking Spike (**Fig. 6**), MLEC can promote CoV replication independent of Spike production.

In parallel, MLEC KD might handicap the glycoprotein quality control network and lead to accumulation of misfolded diglucosylated proteins, thereby perturbing ER proteostasis. CoVs rely on the host ER for infection, from building DMVs from ER membranes to using host ER chaperones (many of which are glycosylated) to fold a high influx of nascent viral proteins. Imbalances in ER proteostasis would lead to a detrimental environment for viral replication and indeed previous work has shown that potent ER stressors lead to hampered CoV infection(Shaban et al., 2021). Our proteomics data reveals that there is only a modest increase in the Heat Shock Factor 1 (HSF1) pathway, while the Unfolded Protein Response is relatively unchanged (**Fig. S15D**). In addition, there are only minor increases in endogenous glycopeptide levels (**Fig. S15E-G**). Together, these results indicate that while MLEC KD leads to some alterations in ER proteostasis and host glycosylation, these changes are modest and may not be the primary mechanism by which MLEC KD hinders CoV replication.

Additional work is needed to characterize the spatial organization of the nsp2-OST-MLEC interaction complex during infection and answer whether MLEC and OST complexes localize to RTCs. MLEC localization has been shown to be regulated by its association with RPN1, where under basal conditions MLEC localizes to the ER but under ER stress conditions can traffic to the Golgi (Yang et al., 2018). RPN1 overexpression restores MLEC localization to the ER. The FT-MLEC interactomics shows that during MHV infection, MLEC retains its interactions with RPN1 and other OST interactors. Additionally, we showed that nsp2 interacts with RPN1 during infection. This suggests that nsp2 may use RPN1 to retain MLEC in the ER to assist in viral protein production. Additionally, MLEC has also been shown to localize to ER-mitochondria contact sites (MAMs)(Carreras-Sureda et al., 2019), which regulate mitochondrial bioenergetics. We have previously shown that SARS-CoV-2 nsp2 and nsp4 can partially localize to MAMs(Davies et al., 2020), so these viral proteins may also dysregulate MLEC and MAMs activity to promote infection.

It will also be important to identify the structural determinants of these interactions, including resolving the binding interfaces by creating truncations and point mutations, such as abolishing the lectin binding residues in MLEC(Takeda et al., 2014). Profiling the kinetics of viral glycoprotein production in the absence of MLEC and developing targeted MS workflows to detect nsp glycopeptides will also aid our understanding of the role of MLEC in viral glycoprotein biogenesis. Lastly, the dependency of MLEC for infection will need to be tested for other RNA viruses.

To our knowledge, this is the first time MLEC has been implicated in CoV infection. Our comparative interactomics approach suggests MLEC plays a role in multiple CoVs, and we show MLEC promotes both MHV and SARS-CoV-2 replication. Prior work has also indicated a role for MLEC in influenza protein folding(Galli et al., 2011), indicating that MLEC may serve a pro-viral function in various RNA virus infections and constitute an attractive broad-spectrum anti-viral target. Indeed, the strategy of targeting host proteostasis factors to inhibit RNA virus replication is actively pursued (Almasy et al., 2021a; Aviner et al., 2023, 2021; Echavarría-Consuegra et al., 2021; Heaton et al., 2016; Sims et al., 2021; Taguwa et al., 2019). Additionally, MLEC remains an understudied component in glycoprotein biogenesis. Our rich interactomics data sets, in which we detect interactions with galectins and other glycoprotein processing factors, may shed further light on the role of MLEC within the glycoprotein biogenesis pathway. In summary, our work combines quantitative proteomics with classical virology to identify the conserved nsp2 interactor MLEC and reveal its pro-viral function in mediating nsp protein biogenesis and CoV replication.

## Materials and Methods

### Plasmids

Expression constructs for SARS-CoV-2, SARS-CoV, and hCoV-OC43 nsp2 and nsp4 homologs were used as previously reported(Davies et al., 2020). Expression sequences for nsp2 and nsp4 homologs of MERS-CoV, hCoV-2293, and MHV were pulled from GenBank (NC_019843 MERS-CoV EMC/2012; AF304460 hCoV-229E; KF268337 MHV-icA549). For nsp4, a corresponding 18 (MHV), 23 (MERS) or 21 (229E) aa leader sequence from the C-terminus of nsp3 was added to the N-terminus, as performed previously (Kanjanahaluethai et al., 2007; Oudshoorn et al., 2017). Sequences were human optimized, synthesized, and cloned into a pTwist-CMV-Hygro vector with an N-terminal 3x-FLAG tag (nsp2) or C-terminal 3x-FLAG tag (nsp4) (Twist Biosciences).

Expression construct for FT-MLEC was made using the mouse coding sequence (GenBank, NP_780612.2). The endogenous signal sequence (1-29 aa) was replaced with the pro-trypsin signal sequence to improve expression efficiency (MSALLILALVGAAVA), followed by a 3x-FLAG tag prior to the remaining MLEC sequence. Construct was mouse codon optimized, synthesized and cloned into a pTwist-CMV-Hygro vector (Twist Biosciences).

SARS-CoV-2 replicon plasmids for Delta and Omicron variants were provided by Drs. Melanie Ott and Taha Taha (Gladstone Institutes). Replicons are encoded on a pBAC vector with CMV and T7 promoters(Taha et al., 2023). GFP encoded on a pcDNA3.1 backbone was used as a transfection control.

### Cell Lines

HEK293T cells were a generous gift from Dr. Joseph Genereux (University of California, Riverside) and grown in high glucose Dulbecco’s Modified Eagle Growth Medium with 10% fetal bovine serum, 1% glutamine, and 1% penicillin/streptomycin (DMEM-10). Murine Delayed Brain Tumor (DBT) cells were a generous gift from Dr. Mark Denison (Vanderbilt University Medical Center) and grown in DMEM-10 with 10 mM HEPES (DMEM-10-HEPES). HEK293T and DBT cells were maintained at 37°C with 5% CO2 and passaged with 0.25% (HEK293T) or 0.05% (DBT) Trypsin-EDTA. Cells were tested regularly for mycoplasma.

To generate stable expressing FT-MLEC DBT cells, parental DBT cells were transfected with 1.5 µg pTwist-CMV-Hygro-FT-MLEC plasmid in 6-well plates using 4.5 uL FuGENE 4K transfection reagent (#E5911) in DMEM-10-HEPES media (antibiotic-free) according to the manufacturer’s directions. After two days, cells were expanded into 10 cm plates and selected with 200 µg/mL Hygromycin in DMEM-10-HEPES media. After an additional three days, cells were selected with 100 µg/mL Hygromycin in DMEM-10-HEPES media for another eleven days, with periodic media changes. Surviving polyclonal cells were expanded and frozen down, with a portion of cells lysed and tested for FT-MLEC expression by Western blot.

### Virus Stocks

MHV-A59 (MHV-WT), MHV-FFL2 (Firefly luciferase fused to the N-terminus of nsp2)(Freeman et al., 2014), MHV-GFP2 (GFP fused to the N-terminus of nsp2)(Freeman et al., 2014), and MHV-del2 (deletion of nsp2)(Graham et al., 2005) were also gifted from Dr. Mark Denison (Vanderbilt University Medical Center). Virus was propagated in 10 cm dishes of DBT cells by infecting at MOI 0.01 for 24 hpi in DMEM-2-HEPES (same recipe as DMEM-10-HEPES, with only 2% FBS), freezing plates at -80°C, then thawing, centrifuging media (4,000 rpm, 10 min, 4°C), and storing supernatant in -80°C. Low passage stocks were used for infection experiments.

### Transient Transfection

HEK293T cells were seeded into 10 cm dishes at 1.5 x 10^6^ cells/plate in DMEM-10 media and a day later transfected with 5 µg of respective nsp plasmids or Tdtomato control via drop-wise addition of a calcium phosphate mixture (0.25 M CaCl_2_, 1x HBS (137 mM NaCl, 5 mM KCl, 0.7 mM Na_2_HPO_4,_ 7.5 mM D-glucose, 21 mM HEPES)). Media was exchanged for fresh DMEM-10 16 h post-transfection, and cells were harvested by scraping an additional 24 h later using cold PBS (containing 1 mM EDTA) on ice, followed by x2 PBS washes. Cells were lysed in TNI lysis buffer (50mM Tris pH 7.5, 150mM NaCl, 0.5% IGEPAL-CA-630) containing 1x Roche c0mplete EDTA-free protease inhibitor (#4693132001) on ice for 15 min, sonicated 10 min at room temperature (RT), and then spun at 21,100xg (15 min, 4°C) to clear lysates of cell debris. Supernatants were transferred to new tubes. Protein concentration was measured by Pierce BCA kit (Thermo Fisher Scientific, #23225) and normalized.

### SDS-PAGE and Western Blotting Analysis

Samples were combined with a 6x Laemelli buffer (12% SDS, 125 mM Tris, pH 6.8, 20% glycerol, bromophenol blue, 100 mM DTT) at a 5:1 ratio, then heated at 37°C for 30 min. Samples were run on an SDS-PAGE gel before being transferred to Immobilon-FL PVDF membrane paper (Millipore Sigma, 0.45 um, #IPFL00010). Blots were blocked with 5% milk in TBS-T (Tris-buffered saline with 0.1% Tween) for 1 h, RT then probed with primary antibody. Anti-M2 FLAG (Sigma Aldrich, #F1804), anti-M1 FLAG (Sigma Aldrich, #F3040), anti-GAPDH (Genetex, #GTX627408), anti-ERDJ3 (Santa Cruz Biotechnology, #SC-271240), or anti-GFP (produced in-house) at 1:1000 in blotting buffer (5% BSA with 0.01% Sodium Azide in TBS-T). Anti-MLEC antibody (ProteinTech, # 26655-1-AP) was used at 1:800 in blotting buffer. Blots were then treated with either 1:10,000 dilutions in 5% milk/TBS-T of goat anti-mouse IgG StarBright700 (BioRad, #12004158) or goat anti-rabbit IgG IRDye800CW (Fischer, #926-32211) secondary antibody or 1:10,000 dilutions in TBS-T of anti-mouse or anti-rabbit IgG HRP (Promega, #W4021 or #W4011). All blots were imaged by ChemiDoc and analyzed on BioRad ImageLab.

### siRNA Reverse Transfection

siGENOME SMARTpool or individual mouse siRNAs for interactors, siGENOME Non-targeting Control (Scramble; # D-001206-13), and siTOX (#D-001500-01-05) were purchased from Horizon. In 96-well plate experiments, plates were pre-treated with poly-D-lysine (0.1 mg/mL) for 1 min at 37°C and washed twice with PBS. Experiments in 12-well and 6-well dishes were not pre-treated with poly-D-lysine. DBT cells (1x10^4^, 96-well; 7.5x10^4^, 12-well; 1.5x10^5^, 6-well) were reverse transfected with siRNA (25 nM final concentration) using 1% DharmaFECT 1 Transfection Reagent (Horizon/Dharmacon, #T-2001-02) according to the manufacturer’s instructions. Opti-MEM was used to dilute siRNAs and DharmaFECT 1 transfection reagent, with antibiotic-free DMEM-10-HEPES media used to culture cells. Transfected cells were then incubated at 37°C for 40 h, prior to MHV infection.

### CellTiter-Glo Assay

DBT cells were seeded in 96-well plates and reverse transfected with siRNAs in triplicate as detailed above. After 40 h, media was removed, cells washed with PBS, and analyzed by Cell TiterGlo kit (Promega, #G7572) according to the manufacturer’s instructions.

### MHV Infection

Viral stocks were thawed on ice prior to use. DBT cells were counted, and the amount of virus added to cells was calculated based on respective MOI. Virus stock was mixed thoroughly with 25 µL (96-well), 100 µL (12-well), or 200 µL of DMEM10-HEPES or antibiotic-free DMEM-10-HEPES (for siRNA transfected samples) per well. Cells were washed with PBS, then inoculum added, and plates incubated for 30 min at 37°C, with rocking every 10 min. Inoculum was removed and cells washed twice with PBS before addition of respective media. Cells were incubated at 37°C for the prescribed infection duration. At harvest, supernatant was removed and saved for plaque assay. Cells were washed with PBS and then harvested by 0.05% Trypsin-EDTA.

### Plaque Assay

DBT cells were plated in technical duplicate/dilution at 2.5x10^5^ cells/well in 12-well plates. After 24 h, serial 1:10 dilutions of virus samples were made in gel saline (3 mg/mL gelatin from porcine skin (Sigma Aldrich, #G2625) in DPBS (Gibco, #14040-133). 50 µL of respective viral dilutions were added to cells and incubated for 30 min at 37°C (rocking every 10 min). After, 1 mL of overlay solution of 1:1 2% agar:2xDMEM-10-HEPES was added to wells and allowed to solidify (5 min, RT), before plates were incubated at 37°C for 20 hpi or until visible plaques formed. The assay was stopped with addition of 4% formaldehyde in PBS (20 min, RT) and solid overlay removed. Plates were air dried, and plaques counted manually.

### MHV-FFL2 Replicase Reporter Assay

DBT cells were infected with MHV-FFL2 as detailed above. At harvest, supernatant was removed, and cells washed with PBS. Assays in 96-well plates (RNAi interactor screen) were then lysed directly with 100 µL/well of SteadyGlo substrate (Promega, # E2510) and 5 min of low rpm shaking. Assays in 12-well plates (all others) were lysed with 200 µL of Glo Lysis Buffer (Promega, #E2661) for 5 min (RT), spun at 13,000xg, 2 min (RT), and mixed in a 1:1 ratio with SteadyGlo substrate in three technical replicates/sample in a 96-well plate. Plates were measured for luminescence using a BioTek Synergy LX Multimode plate reader at 255 gain. A portion of lysates was retained for Western blot analysis.

### SARS-CoV-2 Replicon Assay

HEK293T cells were seeded at 8x10^4^ cells/well in 12-well plates in antibiotic-free DMEM-10-HEPES. Twenty-four hours later, cells were transfected with 500 ng of SARS-CoV-2 Delta or Omicron replicon plasmid and 25 nM (final concentration) human siMLEC siRNA (Horizon, siGENOME #M-020316-00-0005). GFP plasmid (500 ng) and Scramble siRNA (Horizon, siGENOME Non-targeting Control, # D-001206-13) were used as controls. DNA and siRNA were mixed with 3% DharmaFECT Duo (Dharmacon, # T-2010-02) in up to 100 µL of Opti-MEM and incubated at room temperature for 20 min according to the manufacturer’s instructions, before mixing with 800 µL antibiotic-free DMEM-10-HEPES and addition to cells. After 72 h, supernatant was collected, and cells harvested via trypsin.

To measure secreted nanoGlo luciferase reporter in the supernatant, 100 µL of supernatant was mixed with 100 µL of nanoGlo luciferase working reagent (Promega, #N1110) in triplicate, shaken on an orbital shaker for 3 min at room temperature, and luminescence measured as described in “MHV-FFL2 Replicase Reporter Assay”. To measure internal luciferase reporter, cell pellets were lysed in 60 µL cold RIPA lysis buffer (50 mM Tris pH 7.5, 150 mM NaCl, 0.1% SDS, 1% Triton X-100, 0.5% deoxycholate) with 1x Roche c0mplete EDTA-free protease inhibitor (#4693132001) for 15 min on ice, then vortexed twice for 10 s each. Lysates were clarified with a 21,100xg spin at 4°C for 15 min and transferred to fresh tubes. Then 10 µL of sample was mixed with 100 µL nanoGlo luciferase working reagent in duplicate and luminescence measured as detailed above. Cell lysates were also analyzed by Western blot, as detailed in “SDS-PAGE and Western Blotting Analysis”.

### Immunofluorescence

FT-MLEC DBT cells were plated on glass-bottom culture dishes (MatTek, #P35G-0-14-C) and grown for 24 h. Cells were fixed with 4% paraformaldehyde in PBS, washed x3 with PBS, then permeabilized in 0.2% Triton-X/PBS. After three PBS washes, cells were treated with blocking buffer (1% BSA/PBS with 0.1% Saponin). After blocking, cells were incubated with a 1:750 dilution of anti-PDIA4 primary antibody (Protein Tech, 14712-1-AP) in blocking buffer for 1 hour at 37°C. After three PBS washes, cells were incubated with AlexFluor 488-conjugated anti-rabbit goat antibody (ThermoFisher, A-11008) in blocking buffer (1:500 dilution) for 30 min at RT. Cells were then stained with 1:1000 dilution of M2 FLAG primary antibody (SigmaAldrich, F1804) and 1:500 AlexFluor 594-conjugated anti-mouse goat antibody (ThermoFisher, A-11005) in blocking buffer. Cells were then mounted in Prolong Gold with DAPI stain (ThermoFisher, P36935). Cells were imaged using a Zeiss LSM-880 confocal microscope and analyzed in ImageJ.

### Quantitative Reverse Transcriptase Polymerase Chain Reaction (RT-qPCR)

Cellular RNA was extracted with a QuickRNA miniprep kit (Zymo, #R1055). Then, 500 ng total cellular RNA was reverse transcribed into cDNA using M-MLV reverse transcriptase (Promega, #M1701) with random hexamer primers (IDT, #51-01-18-25) and oligo-dT 15mer primers (IDT, #51-01-15-05). For negative-sense gRNA measurement, primer #3 (**Table 1**) was used instead to generate cDNA. iTaq Universal SYBR Green Supermix (BioRad, #1725120) was then used with respective primers (primers #1-2, 4-11) for target genes and reactions run in technical duplicate in 96-well plates on a BioRad CFX qPCR instrument. Amplification was carried out in the following sequence: 95°C, 2 min; 45 cycles of 95°C, 10 s and 60°C, 30 s. A melting curve was generated in 0.5°C intervals from 65°C to 95°C. Cq values were calculated in the BioRad CFX Maestro software and transcripts were normalized to a housekeeping gene (*Gapdh* or *Rpl13a*).

### NGI-1 Treatment

To determine dosage, DBT cells were seeded in 12-well plates, then 4 h later treated with a range of doses (0.05, 0.5, 5, 10, 20 µM) of NGI-1 (SigmaAldrich, #SML1620), Tunicamycin (Tm, 1 µg/mL; Abcam, #ab120296), or DMSO. After 20 h, cells were harvested and lysed in TNI lysis buffer as described in “Transient Transfection.” DMSO-treated lysates were split in half, with one half deglycosylated by PNGase F (Promega, #V4831) treatment for 1 h at 37°C. Lysates were then analyzed by Western blot.

To test MLEC KD and NGI-1 effects on MHV-FFL2, 40 h post-siRNA reverse transfection DBT cells were infected MHV-FFL2 as described above. After inoculum incubation and PBS washes, media containing 5 µM NGI-1 or DMSO was added to cells. The 5 uM NGI-1 dosage was chosen as it resulted in partial inhibition of glycosylation while not completely blocking MHV infection. Infected cells were measured for luminescence as detailed above.

### FLAG Immunoprecipitations (IPs)

Sepharose 4B resin (SigmaAldrich, #4B200) and G1 anti-DYKDDDDK resin (GenScript, #L00432-10) were pre-washed four times with wash buffer (lysis buffer without protease inhibitor) for each sample. Cell lysates were normalized in wash buffer and added to 15 µL Sepharose 4B resin for 1 h, rocking at 4°C. Resin was collected by centrifugation for 10 minutes at 400xg (4°C) and pre-cleared supernatant was added directly to 15 µL G1 anti-DYKDDDDK resin and rocked at 4°C overnight. Supernatant was then removed, and resin was washed x4 with wash buffer. Bound proteins were eluted with elution buffer (62.5 mM Tris, pH 6.8, 6% SDS) for 30 min at RT then 15 min at 37°C, followed by a second elution for 15 minutes at 37°C.

### GFP IP

DBT cells were infected with MHV-GFP2 or MHV-WT in 10 cm plates (MOI 1, 9 hpi) as detailed above. Cells were harvested by 0.05% trypsin-EDTA and lysed in 200 µL TNI lysis buffer with protease inhibitor (Roche, #4693132001) as detailed in “Transient Transcription.” Normalized lysates were pre-cleared with 15 µL Sepharose 4B resin as in “FLAG IPs” and then cleared supernatant was added to 15 µL GFP-Trap magnetic agarose beads (Chromotek, #GTMA-20) and incubated for 2 h rocking (4°C). Beads were separated with a magnet and rinsed four times with wash buffer. Proteins were eluted with elution buffer as in “FLAG IPs.”

### FT-MLEC IP and DSP Crosslinking

Parental or FT-MLEC DBT cells were infected with MHV (MOI 2.5, 9 hpi) in 10 cm plates as detailed above. At harvest, cells were washed x3 with PBS, then incubated with 10 mL of 0.5 mM dithiobis(succinimidyl propionate) (DSP) crosslinker (ThermoScientific, #PI22585) in PBS at 37°C for 10 min. Crosslinker was quenched with 1 mL of 1 M Tris (pH 7.5) for 5 min at 37°C, after which cells were washed x3 with PBS and trypsinized to remove from plate. Cells were lysed in RIPA lysis buffer (50 mM Tris pH 7.5, 150 mM NaCl, 0.1% SDS, 1% Triton X-100, 0.5% deoxycholate) with protease inhibitor for 15 min on ice, followed by two 10 sec vortex pulses and spun at 21,100xg (15 min, 4°C). Cleared supernatant was normalized and immunoprecipitated for FLAG as in “FLAG IP.”

### Global Viral Proteomics Samples

DBT cells were reverse transfected with siRNA in 6-well plates and after 40 h infected with MHV (MOI 1, 10 hpi). Cells were harvested by 0.05% trypsin-EDTA and then lysed in TNI lysis buffer and prepared for mass spectrometry as follows.

### General Mass Spectrometry Sample Preparation

Elutions or 20 µg of protein/lysate were prepared for mass spectrometry. Proteins were precipitated with LC/MS-grade methanol:chloroform:water (in a 3:1:3 ratio), then washed three times with methanol, with each wash followed by a 2-minute spin at 10,000xg at RT. Protein pellets were air dried and resuspended in 5 µL of 1% Rapigest SF (Waters, #186002122). Resuspended proteins were diluted with 32.5 µL water and 10 µL 0.5 M HEPES (pH 8.0), then reduced with 0.5 µL of fresh 0.5 M TCEP for 30 min at RT. Samples were alkylated with 1 µL of fresh 0.5 M iodoacetamide for 30 min in the dark at RT and then digested with 0.5 µg Pierce Trypsin/Lys-C (Thermo Scientific, # A40007) overnight at 37°C shaking. Peptides were diluted to 60 µL with LC/MS-grade water and labeled using TMTpro labels (Thermo Scientific, # A44520) for 1 h at RT, followed by quenching with fresh ammonium bicarbonate (0.4% v/v final) for 1 h at RT. Samples were then pooled, acidified to pH < 2.0 with formic acid, concentrated to 1/6^th^ original volume via Speed-vac, and diluted back to the original volume with buffer A (95% water, 5% acetonitrile, 0.1% formic acid). Cleaved Rapigest products were removed by centrifugation at 17,000xg for 30 min (RT) and supernatant transferred to fresh tubes.

### PNGase F Mass Spectrometry Sample Preparation

DBT cells were seeded in 12-well plates and reverse transfected with siRNA as previously described, then infected with MHV (MOI 1) after 40 h. At 9 hpi, cells were harvested by 0.05% trypsin-EDTA, washed twice with PBS, then lysed in RIPA lysis buffer with protease inhibitor. Protein concentrations were measured by BCA kit and 20 µg protein/sample was carried forward to MS preparation on a BioMek i7 automated workstation. Proteins were reduced with 5 mM DTT (60°C, 30 min, 1,000 rpm shaking), followed by alkylation with 20 mM Iodoacetamide (RT, 30 min, in dark) and quenched with additional 5 mM DTT (15 min, RT). Hydrophilic and hydrophobic Sera-Mag Carboxylate-modified SpeedBeads (Cytiva, #09-981-121, #09-981-123) were prepared in a 1:1 ratio and washed x4 with LC/MS-grade water. Eight (8) µL of beads mixture was added to sample to bind proteins and mixed. Pure ethanol (90 µL) was added to samples and incubated 5 min, RT, at 1,000 rpm. Beads were separated by a magnet and washed x3 with 180 µL 80% ethanol. Proteins were then on-bead digested with 0.5 µg Trypsin/LysC (Thermo Scientific, # A40007) in 0.5 M HEPES buffer (pH 8.0) for 10 h, shaking at 1,000 rpm at 37°C. After digestion, samples were labeled with TMTpro labels (1 h, RT) and quenched with 10% w/v ammonium bicarbonate (1 h, RT), then pooled together and reduced to 1/5^th^ volume by Speed-vac. Approximately 40 µg of labeled peptides were then treated with PNGase F (Promega, #V4831) for 2 h at 37°C and taken forward to LC-MS/MS analysis.

### Tandem Mass Spectrometry (LC-MS/MS) Analysis

Triphasic MuDPIT columns(Washburn et al., 2001) were packed with 1.5cm Aqua 5 µm C18 resin (Phenomenex, #04A-4299), 1.5cm Luna 5 µm SCX resin (Phenomenex # 04A-4398), and 1.5cm Aqua 5 µm C18 resin. TMT-labeled samples (20 µg of lysates or 1/3^rd^ of IP elutions) were loaded onto the microcapillaries via a high-pressure chamber. Samples were washed, with buffer A (95% water, 5% acetonitrile, and 0.1% formic acid) for 30 min. MudPIT columns were installed on the LC column switching valve and followed by a 20cm fused silica microcapillary analytical column packed with 3 µm Aqua C18 resin (Phenomenex, # 04A-4311) ending in a laser-pulled tip. Analytical columns were washed with buffer A for 30 min prior to use. Peptides were fractionated online using liquid chromatography (Ultimate 3000 nanoLC system) and then analyzed with an Exploris480 mass spectrometer (Thermo Fisher). MudPIT runs were carried out by 10µL sequential injections of 0, 10, 20, 30, 40, 50, 60, 70, 80, 90, 100% buffer C (500mM ammonium acetate, 94.9% water, 5% acetonitrile, 0.1% formic acid), followed by a final injection of 90% C, 10% buffer B (99.9% acetonitrile, 0.1% formic acid v/v). Each injection was followed by a 130 min gradient using a flow rate of 500nL/min (0-6 min: 2% buffer B, 8 min: 5% B, 100 min: 35% B, 105min: 65% B, 106-113 min: 85% B, 113-130 min: 2% B). ESI was performed directly from the tip of the microcapillary column using a spray voltage of 2.2 kV, an ion transfer tube temperature of 275°C and an RF Lens of 40%. MS1 spectra were collected using a scan range of 400-1600 m/z, 120k resolution, AGC target of 300%, and automatic injection times. Data-dependent MS2 spectra were obtained using a monoisotopic peak selection mode: peptide, including charge state 2-7, TopSpeed method (3s cycle time), isolation window 0.4 m/z, HCD fragmentation using a normalized collision energy of 36%, 45k resolution, AGC target of 200%, automatic (whole cell lysates) or 150 ms (IP elutions) maximum injection times, and a dynamic exclusion (20 ppm window) set to 60s.

### Mass Spectrometry Data Processing

Identification and quantification of peptides were performed in Proteome Discoverer 2.4 (Thermo Fisher). For viral protein interactomics in HEK293T cells, the SwissProt human database (TaxID 9606, released November 23^rd^, 2019; 42,252 entries searched) with FT-nsp2 and nsp4-FT sequences (11 entries) manually added (42,263 total entries searched) was used. For peptides from experiments in murine DBT cells, the SwissProt mouse database (TaxID 10090, released November 23^rd^, 2019; 25,097 entries searched) with MHV-WT (GenBank: KF268337, 25 entries) or MHV-GFP2 sequences (GenBank: KF268337, with addition of eGFP sequence to the N-terminus of nsp2 as previously detailed(Freeman et al., 2014), 25 entries) or the FT-MLEC sequence (1 entry) manually added (25,122 - 25,123 total entries searched) were used. Searches were conducted with Sequest HT using the following parameters: trypsin cleavage (maximum two missed cleavages), minimum peptide length 6 AAs, precursor mass tolerance 20 ppm, fragment mass tolerance 0.02 Da, dynamic modifications of Met oxidation (+15.995 Da), protein N-terminal Met loss (−131.040 Da), and protein N-terminal acetylation (+42.011 Da), static modifications of TMTpro (+304.207 Da) at Lys, and N-termini and Cys carbamidomethylation (+57.021 Da). For PNGaseF-treated samples, an additional dynamic modification of Asn deamidation (+0.984 Da) was used. Peptide IDs were filtered using Percolator with an FDR target of 0.01. Proteins were filtered based on a 0.01 FDR, and protein groups were created according to a strict parsimony principle. TMTpro reporter ions were quantified considering unique and razor peptides, excluding peptides with co-isolation interference greater than 25%. Peptide abundances were normalized based on total peptide amounts in each channel, assuming similar levels of background, or based on bait protein for IPs where noted. Protein quantification used all quantified peptides. Post-search filtering was carried out to include only proteins with two or more identified peptides.

### Resource Availability

Further information and requests for resources and reagents should be directed to and will be fulfilled by the lead contact, Lars Plate. All unique/stable reagents generated in this study are available from the Lead Contact with a completed Materials Transfer Agreement.

### Data Availability

The mass spectrometry proteomics data are deposited to the ProteomeXchange Consortium via the PRIDE partner repository under the accession code PXD049130. All other necessary data are contained within the manuscript or can be shared by the Lead Contact upon request.

## Author Contributions

Conceptualization, J.P.D., L.P.; Investigation, J.P.D.; Writing – Original Draft, J.P.D.; Writing – Review & Editing, J.P.D., L.P.; Visualization, J.P.D., L.P.; Supervision, L.P.; Funding Acquisition, L.P.

## Major Subject Areas

Biochemistry and Chemical Biology, Microbiology and Infectious Disease

## Acknowledgements

We would like to thank Dr. Mark Denison and Xiaotao Lu (Vanderbilt University Medical Center) for their generous gifts of reagents, protocols, and consultation on experimental design. We thank Drs. Melanie Ott and Taha Taha (Gladstone Institutes) for the SARS-CoV-2 replicon plasmids and protocols. We also thank Plate lab members Jake Hermanson and Lea Barny for technical assistance with the BioMek sample preparation. This work was funded by R35GM133552 (National Institute of General Medical Sciences) and Vanderbilt University start-up funds. J.P.D. was supported by T32GM008554 (National Institute of General Medical Sciences).

