## Supplemental Information and Figures for "The glycoprotein quality control factor Malectin promotes coronavirus replication and viral protein biogenesis"

**This PDF file includes:**

**Supplemental Tables**

Table S1. Nsp2 comparative AP-MS of interacting proteins and peptides
Table S2. Nsp4 comparative AP-MS of interacting proteins and peptides
Table S3. MHV GFP-nsp2 AP-MS of interacting proteins and peptides
Table S4. FT-MLEC AP-MS of interacting proteins and peptides during MHV infection
Table S5. Global proteomics of MHV infection in Mlec-KD vs. WT DBT cells
Table S6. Glycoproteomics of MHV infection in Mlec-KD vs. WT DBT cells

**Supplemental Figures**

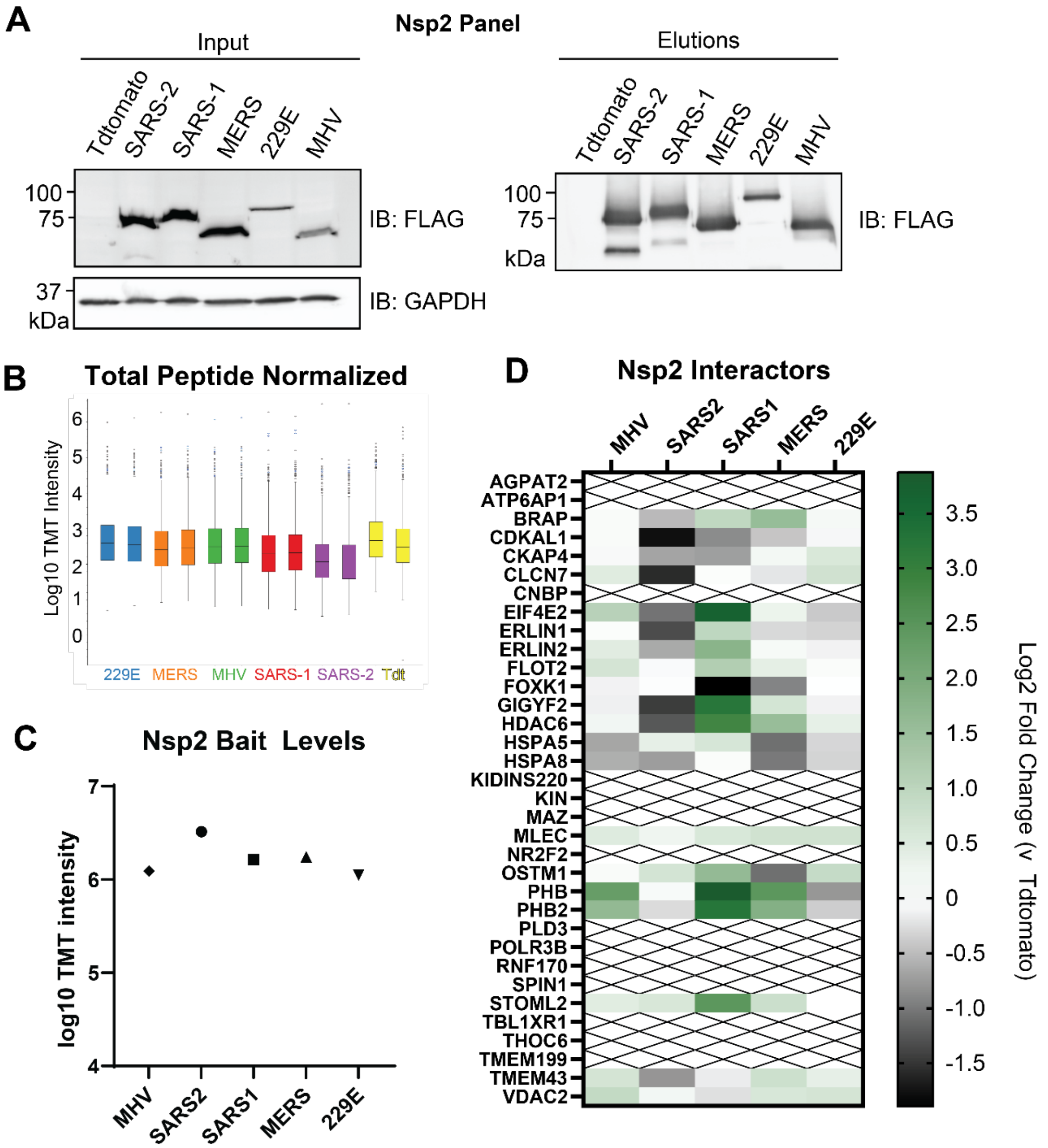

**Figure S1. Nsp2 homolog panel comparative interactome.**

A. Representative Western blot of nsp2 panel from different CoV expressed in HEK293T cells and FLAG immunoprecipitated. Tdtomato serves as negative IP control. Blotting for FLAG and GAPDH as a loading control.
B. Normalized TMT channel abundances of nsp2 panel AP-MS (n = 3-4 IPs/homolog, 1 MS run)
C. Average nsp2 bait levels as measured by AP-MS.
D. Interacting proteins from our previous comparative proteomics study(Davies et al., 2020) were cross-referenced against this nsp2 AP-MS panel and the log2 fold enrichment against Tdtomato background is displayed. Proteins not identified in this the current AP-MS panel are marked with an "X".

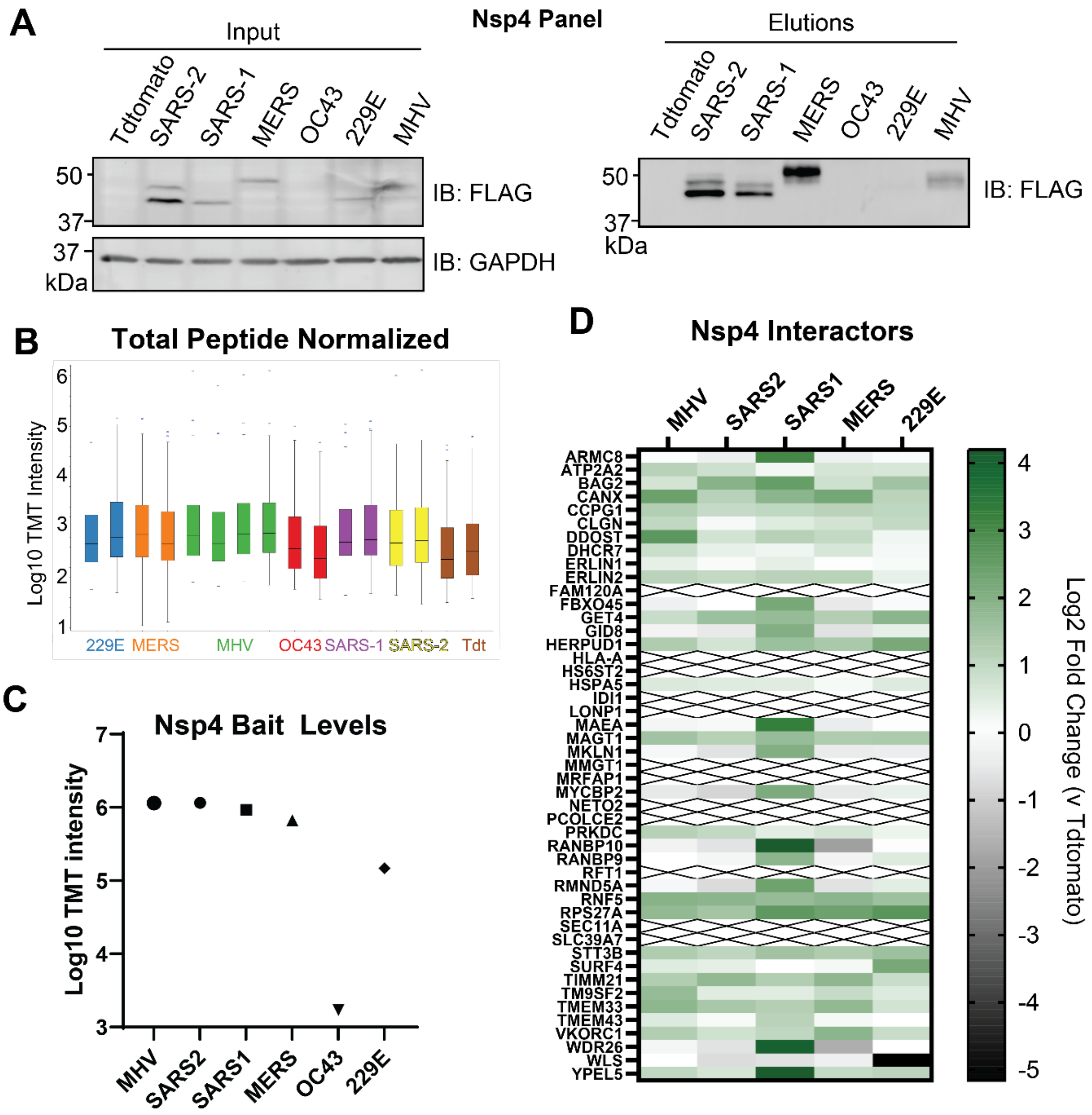

**Figure S2. Nsp4 homolog panel comparative interactome.**

A. Representative western blot of nsp4 homolog panel expressed in HEK293T cells and FLAG immunoprecipitated. Tdtomato serves as negative IP control. Blotting for FLAG and GAPDH as a loading control.
B. Normalized TMT channel abundances of nsp4 panel AP-MS (n = 3-4 IPs/homolog, 1 MS run)
C. Average nsp4 bait levels as measured by AP-MS. OC43 was omitted from analysis due to low expression levels.
D. Interacting proteins from our previous comparative proteomics study(Davies et al., 2020) were cross-referenced against this nsp4 AP-MS panel and the log2 fold enrichment against Tdtomato background is displayed. Proteins not identified in this the current AP-MS panel are marked with an “X”.

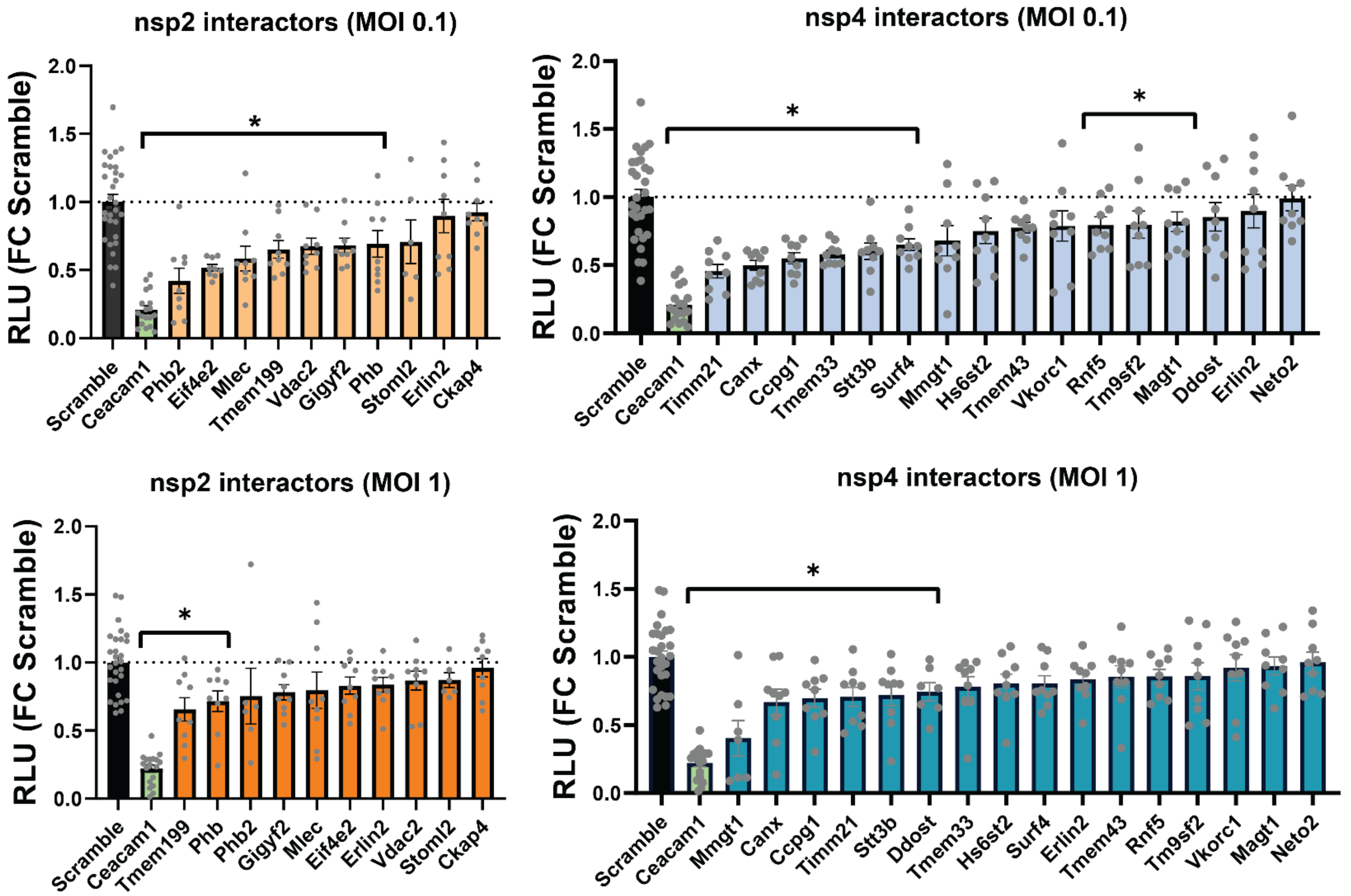

**Figure S3. MHV-FFL2 replicase reporter screen against RNAi of interactors.**

Ten nsp2 interactors and 16 nsp4 interactors were knocked down by siRNA in DBT cells in 96-well plates, then infected with MHV-FFL2 at MOI 0.1 (10 hpi) or MOI 1 (8 hpi). Lysates were harvested and luminescence measured by SteadyGlo assay (Relative Luminescence Units, RLU). Student's T-test for significance, with $p<0.05$ considered significant. KDs that elicit a significant change compared to Scramble are annotated by the asterisk bracket. Mean ±SEM; interactor KD, n = 9 across 3 separate assays; Scramble, n = 28 across 10 separate assays; Ceacam1 KD, n = 18 across 6 separate assays.

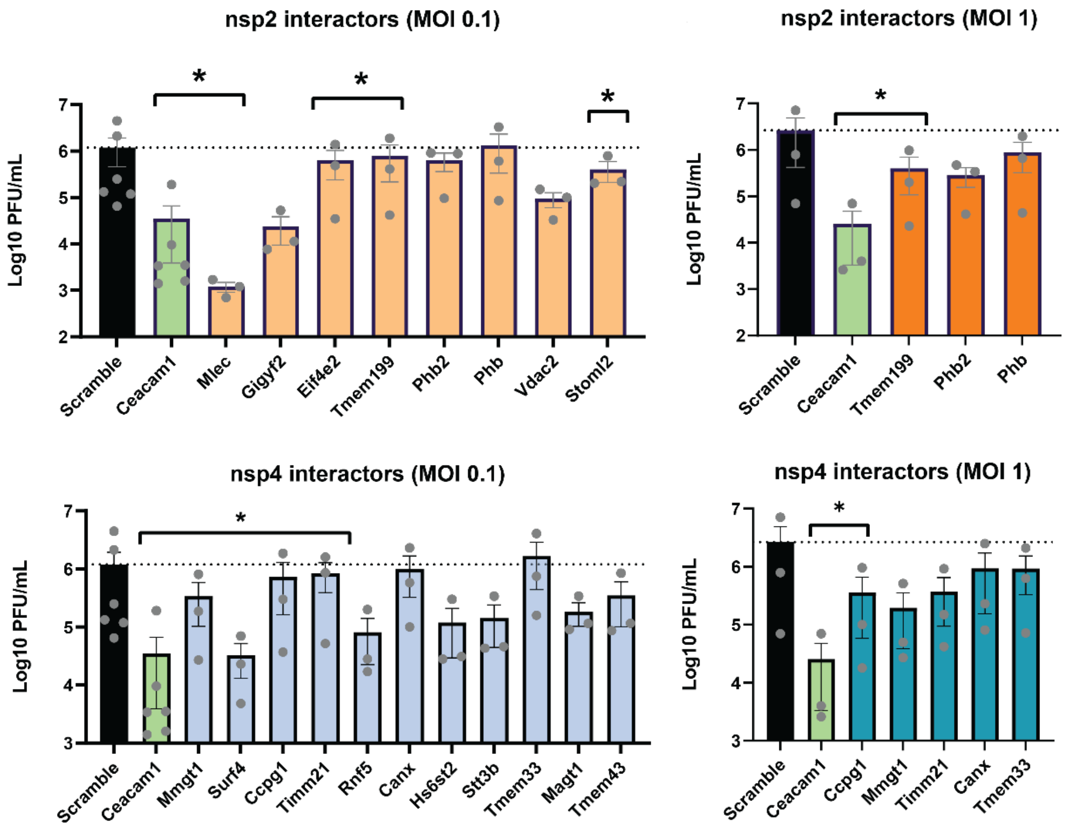

**Figure S4. Effect of interactor knockdown screen on MHV titer.**

Prioritized interactors from the MHV-FFL2 replicase reporter screen were knocked down in DBT cells with siRNA for 40 h, then infected with MHV (MOI 0.1, 10 hpi or MOI 1, 8 hpi as indicated). Scramble siRNA was used as base line control and siCeacam1 as positive control. Supernatant from infected cells was collected and infectious titer measured by plaque assay. Student's t-test for significance, with $p<0.05$ considered significant. Mean±SEM; Interactors, n = 3; Scramble, n = 6 (MOI 0.1) or 3 (MOI 1); siCeacam1, n = 6 (MOI 0.1) or 3 (MOI 1).

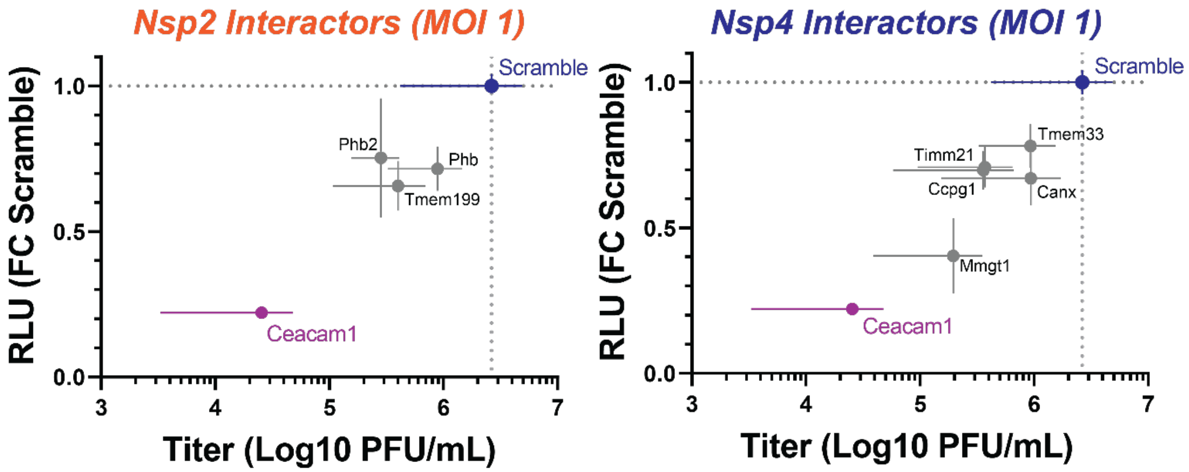

**Figure S5. Effect of interactor knockdown on MHV-FFL2 replicase reporter vs. MHV titer (MOI 1).**

Correlation plot of the effect of prioritized nsp2 and nsp4 interactor knockdowns on MHV-FFL2 replicase reporter vs. MHV titer (MOI 1, 8 hpi). Scramble (blue) siRNA was used as basal control, siCeacam1 (magenta) as a positive control.

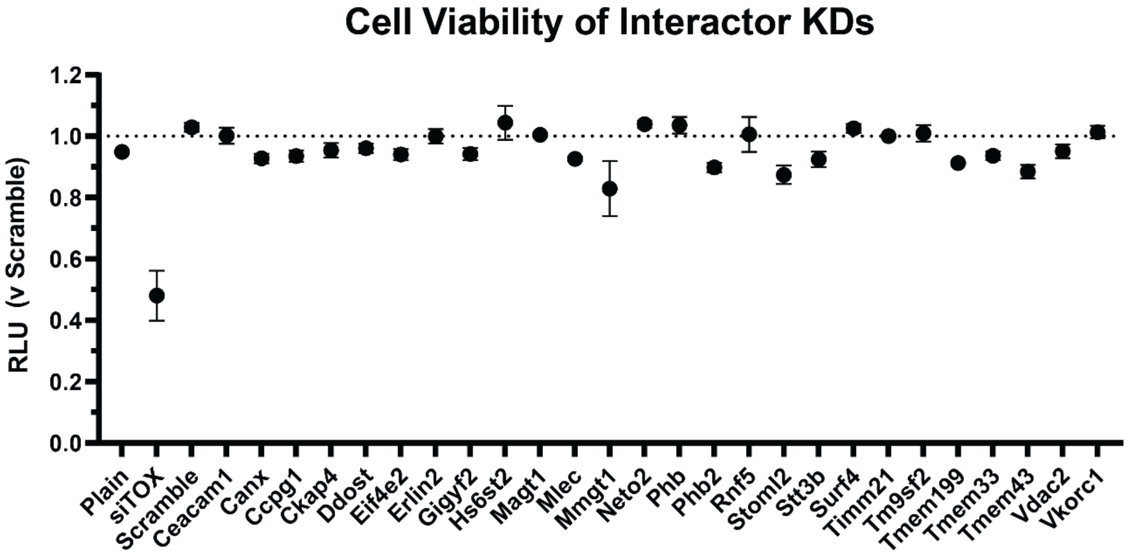

**Figure S6. Cell viability of interactor knockdowns.**

DBT cells were treated with respective siRNAs for 40 hpi in 96-well plate and then assayed for cell viability via CellTiter Glo kit. Non-treated cells (Plain) and siTOX (positive control leading to cell death) were included as controls. Mean±SEM; Scramble, n = 12; siCeacam1, n = 6, all others, n = 3.

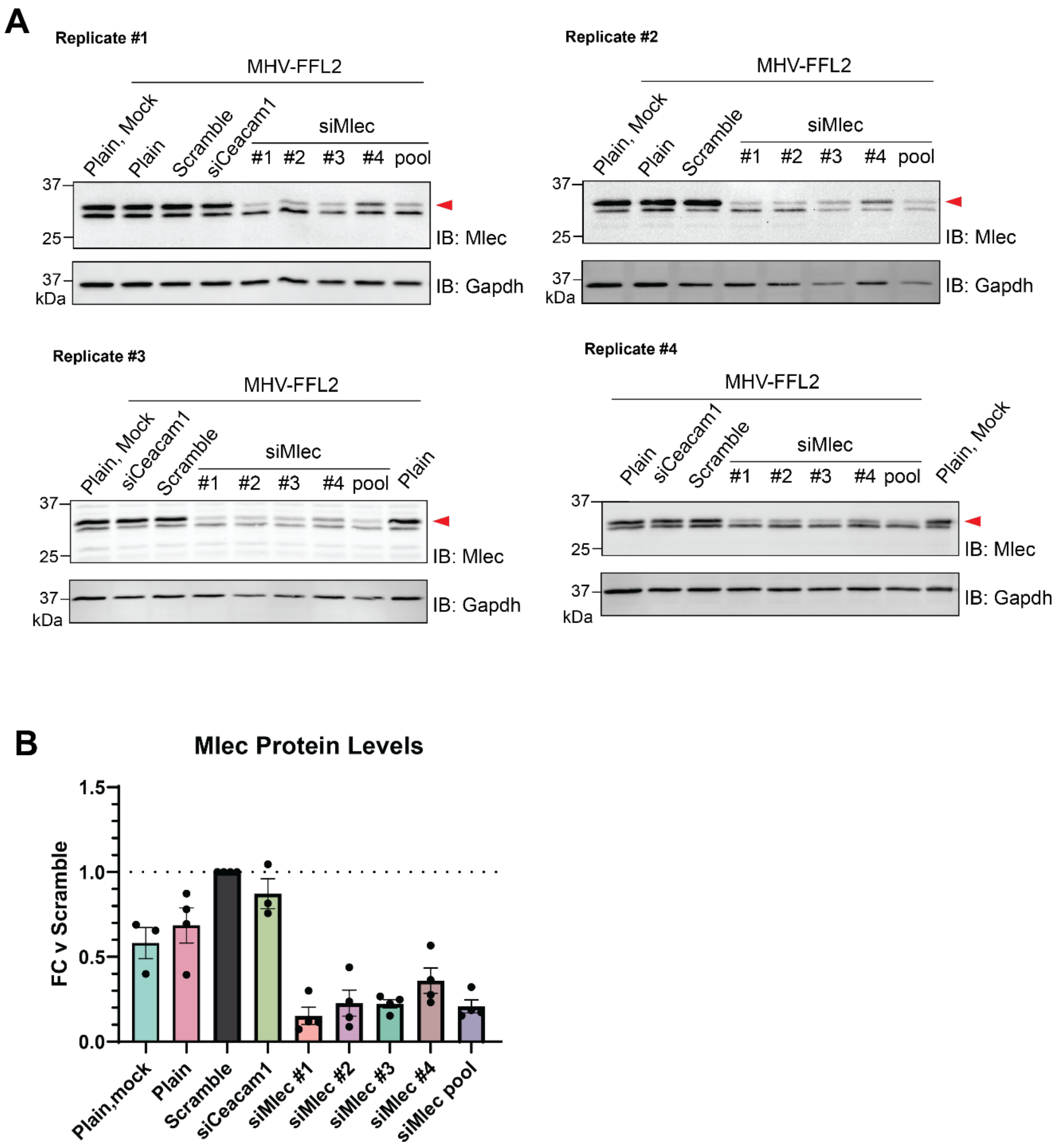

**Figure S7. Mlec KD efficiency with individual vs. pooled siMlec.**

A. Western blots of DBT lysates probed for endogenous Mlec (red arrow) with GAPDH as loading control. Cells were treated with individual siMlec siRNAs (#1-4) or pooled siMlec (all four individual siRNAs combined) for 40 h prior to MHV-FFL2 infection. Plain cells were not treated with siRNA.
B. Quantification of Mlec bands in (A), normalized to GAPDH levels and relative to Scramble control. Mean±SEM, n = 3-4.

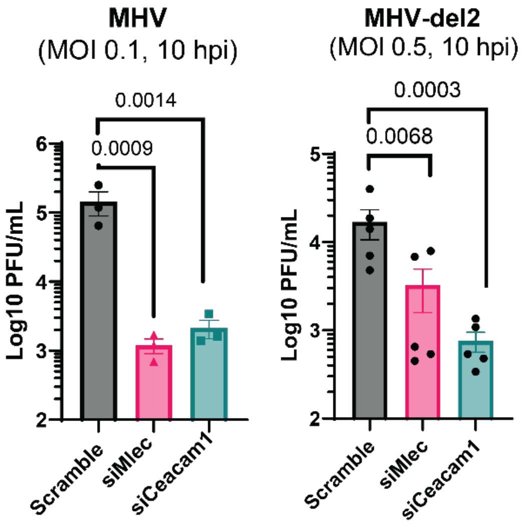

**Figure S8. Genetic interaction between nsp2 and Mlec.**

Infectious titers (PFU/mL), as measured by plaque assay, from DBT cells treated with Scramble, siCeacam1, or siMlec siRNA for 40 h prior to infection with MHV (MOI 0.1, 10 hpi) or MHV-del2 (nsp2 deletion, MOI 0.5, 10 hpi). Mean±SEM; MHV, n = 3; MHV-del2, n = 5. Student's t-test for significance, $p<0.05$ considered significant.

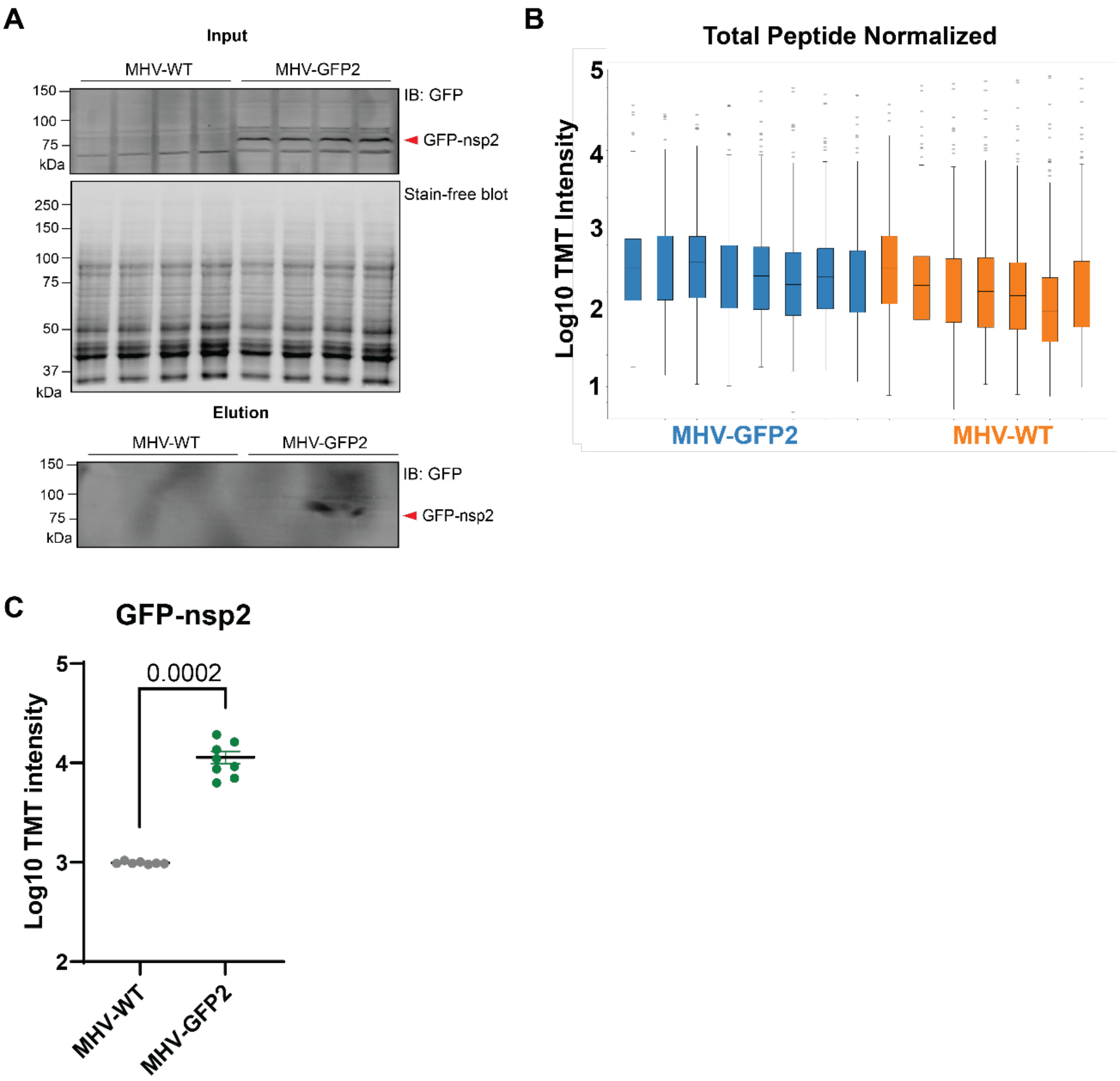

**Figure S9. MHV-GFP2 IP and AP-MS metrics.**

A. Western blots of GFP-IP inputs and elutions for AP-MS (four representative samples per condition), probing for GFP. DBT cells were infected with MHV-WT or MHV-GFP2 (MOI 1, 9 hpi), lysed, and immunoprecipitated for GFP. Stain-free blot of inputs as loading control. Red arrow indicates GFP-nsp2. While we did not detect a GFP-nsp2 band in elution, subsequent MS analysis showed enriched levels of nsp2 over MHV-WT (C), indicating successful enrichment. We attribute lack of blot detection due to poor antibody performance.
B. Total peptide normalized TMT abundance from MHV-GFP2 AP-MS. MHV-WT, n = 7; MHV-GFP2, n = 8 IPs, in 1 MS run.
C. GFP-nsp2 levels measured by AP-MS in MHV-WT and MHV-GFP2 IP samples. One-way ANOVA with multiple testing corrections to test significance.

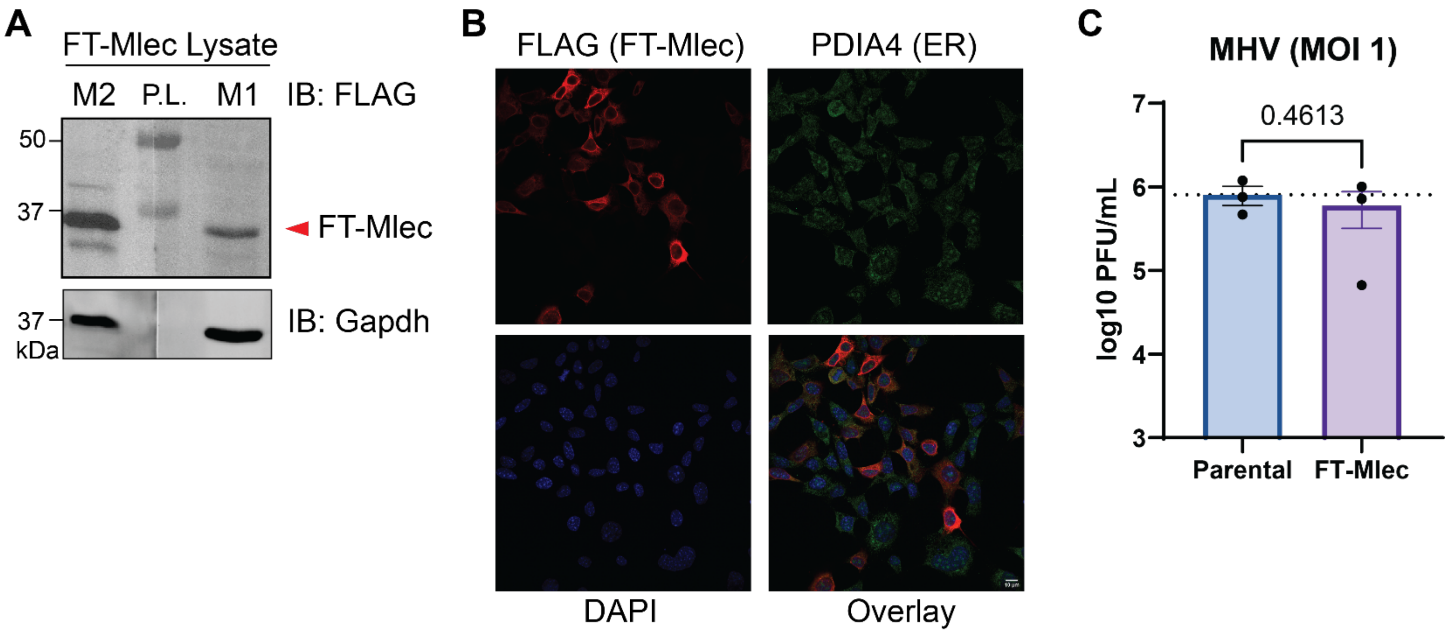

**Figure S10. Validation of FT-MLEC stable DBT cell line.**

A. Western blot of FT-MLEC lysate, probed with either anti-M2 or M1 FLAG, to test for cleavage of the signal sequence and the revealing of the N-terminal FLAG tag (the M1 epitope) on the MLEC construct. The same lysate was split in half and run on an SDS-PAGE gel. After transfer, the blot was cut in two halves and probed separately with respective FLAG antibodies, then imaged together. P.L., protein ladder.
B. Colocalization immunofluorescence (IF) of FT-MLEC DBT cells, staining for FLAG (FT-MLEC), PDIA4 (ER marker) FT-MLEC, and DAPI, with overlay. Scalebar = 10 µm
C. Titers of MHV infection in FT-MLEC or Parental DBT cells (MOI 1, 9 hpi), measured by plaque assay. Paired Student’s t-test for significance, $p<0.05$ considered significant; $n = 3$.

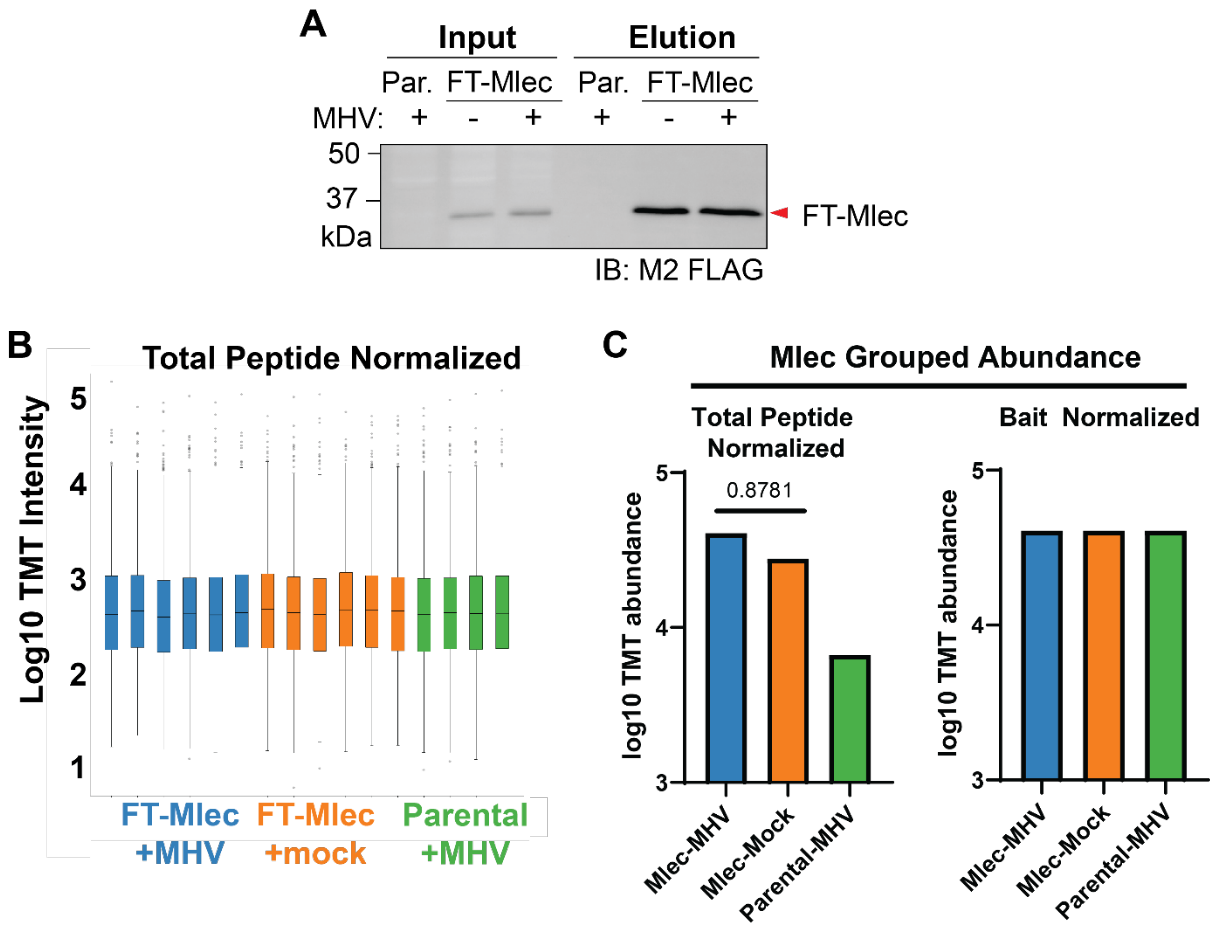

**Figure S11. FT-MLEC IP and AP-MS metrics.**

A. Western blot of representative samples from FLAG IP input and elution in Parental (Par.) or FT-MLEC cells infected with MHV or mock (MOI 2.5, 9 hpi), probed for M2 FLAG. Red arrow indicates FT-MLEC.
B. Total peptide normalized TMT channel abundances of FT-MLEC AP-MS. FT-MLEC + MHV, n = 6; FT-MLEC + mock, n = 6; Parental+MHV, n = 4 IPs.
C. Mlec grouped TMT abundance with total peptide normalization or bait (FT-MLEC) normalization. Bait normalization was used for FT-MLEC interactome comparison between MHV vs. Mock conditions (**Fig. 3F**). T-test with multiple testing corrections for significance.

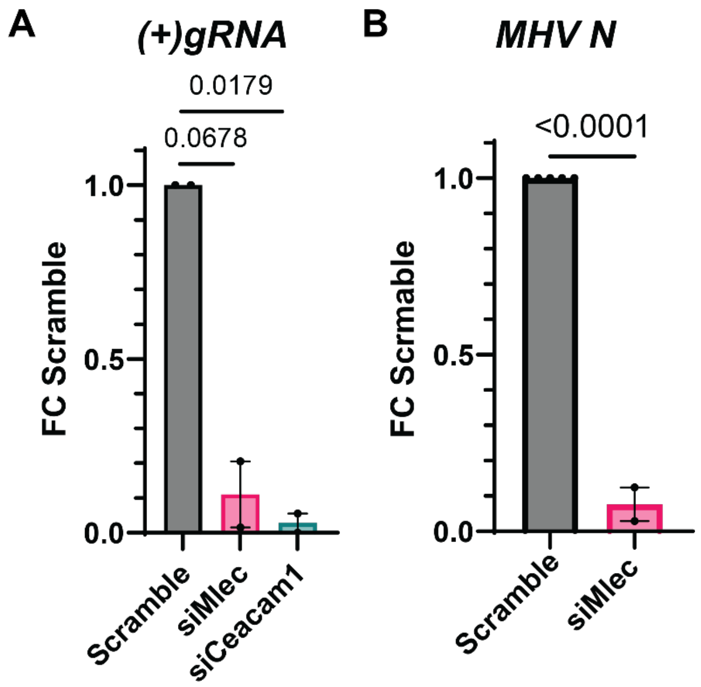

**Figure S12. Effect of Mlec KD on viral genomic and subgenomic RNA levels.**

A. Positive-sense genomic RNA (*(+)gRNA*) levels in Scramble, siCeacam1, or siMlec treated DBT cells infected with MHV (MOI 1, 10 hpi), as measured by RT-qPCR and normalized to *Rpl13a* transcript levels (**Table 1**). n = 2; Student's T-test for significance, $p<0.05$ considered significant.
B. Subgenomic (*Nucleocapsid*) transcript levels in Scramble vs siMlec siRNA treated DBT cells infected with MHV (MOI 0.1, 10 hpi), as measured by RT-qPCR and normalized to *Gapdh* transcript levels (**Table 1**). Scramble, n = 5; siMlec, n = 2; Student's T-test for significance, $p<0.05$ considered significant.

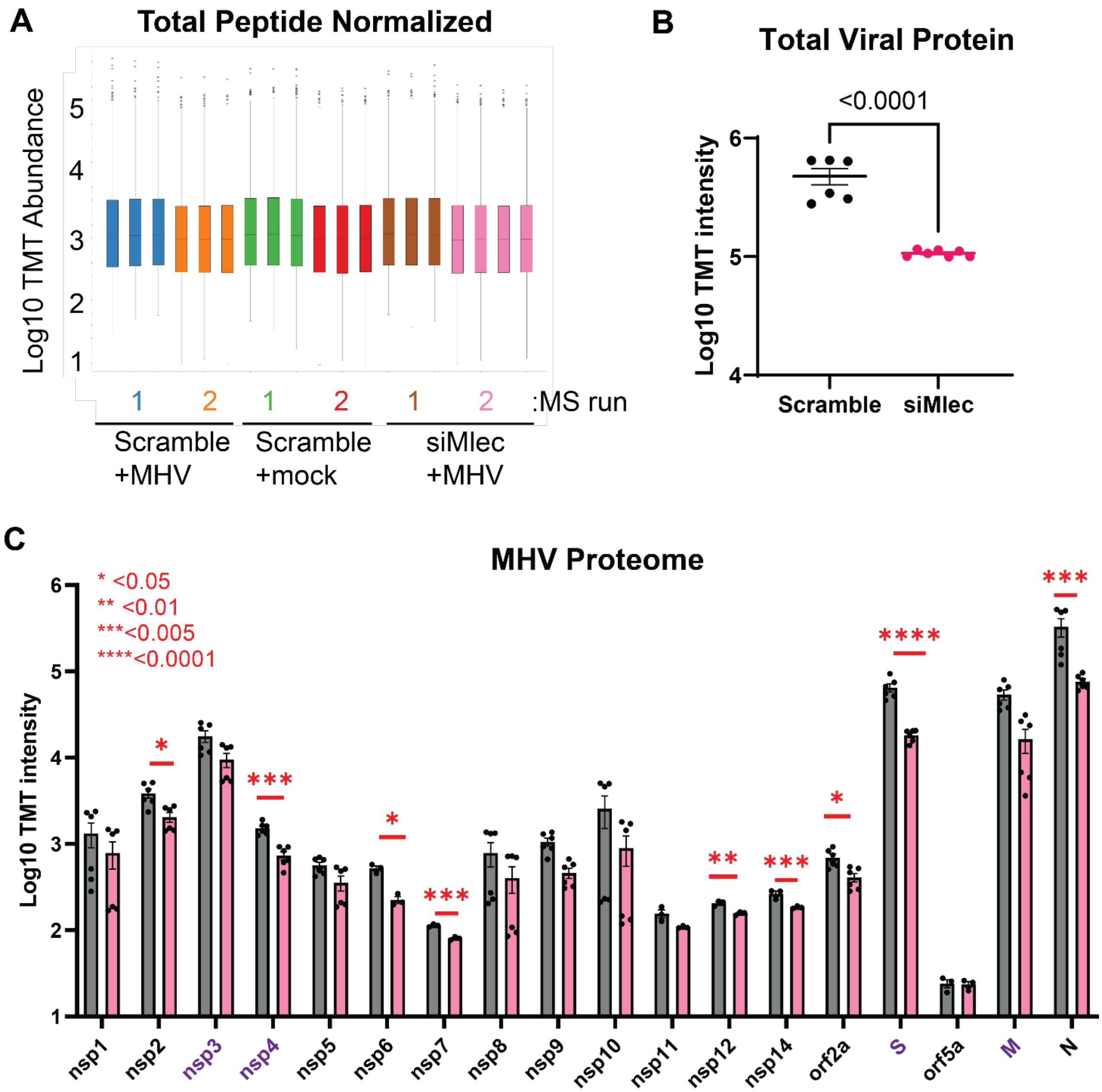

**Figure S13. Effect of Mlec KD on intracellular MHV proteome.**

A. Total normalized TMT channel intensity for global MHV proteome in DBT cells (MOI 1, 10 hpi). Scramble, n = 6; siMlec, n = 7 biological replicates across 2 MS runs.
B. Total viral protein levels in Scramble vs. siMlec treated DBT cells. One-way ANOVA with multiple testing correction.
C. Raw TMT abundance of intracellular viral proteins during MHV infection in DBT cells treated with Scramble or siMlec siRNA (MOI 1, 10 hpi), measured by quantitative proteomics. Purple protein names denote viral *N*-linked glycoproteins. Spike protein (S), membrane protein (M), nucleocapsid protein (N). One-way ANOVA with multiple testing correction. * p<0.05, ** p<0.01, ***p<0.005, ****p<0.0001.

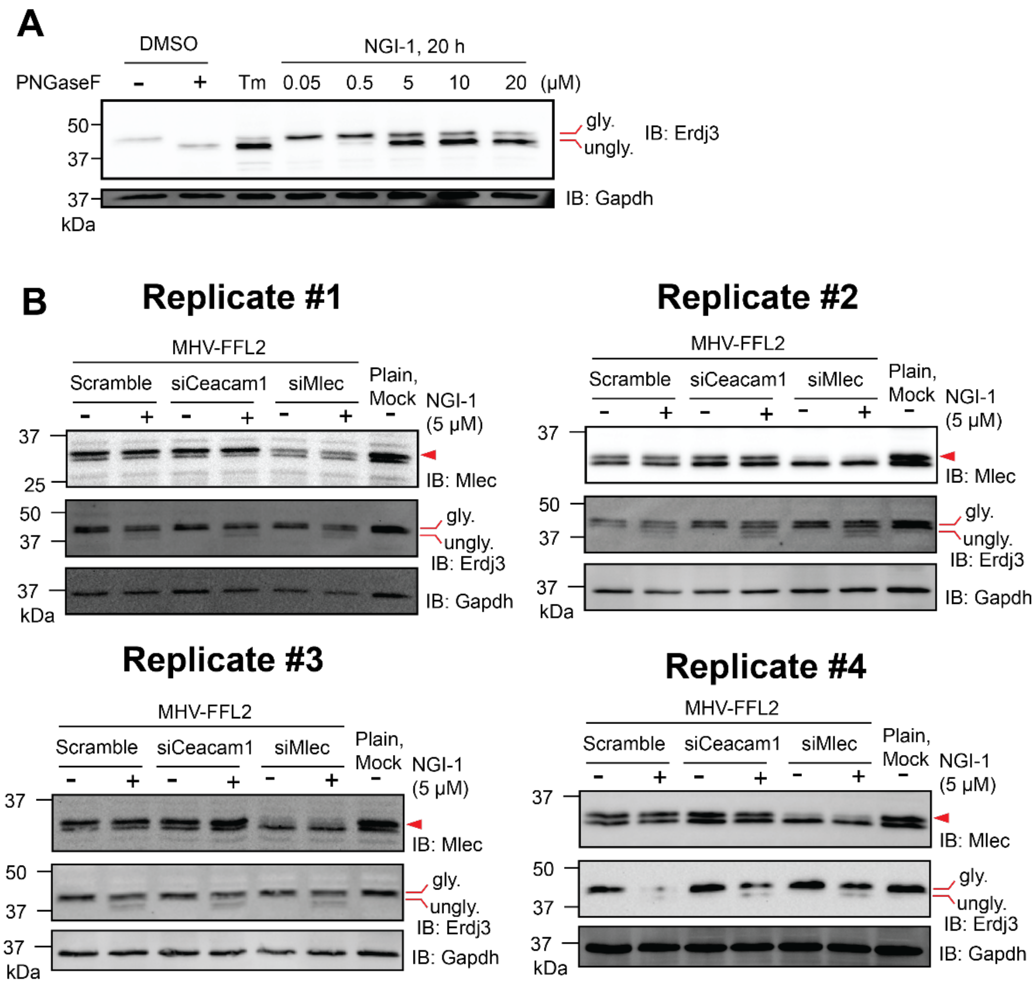

**Figure S14. NGI-1 inhibition of the oligosaccharide transfer complex (OST).**

A. Western blot of DBT cell lysates treated with a range of NGI-1 doses (0.05, 0.5, 5, 10, 20 µM) for 20 h prior to harvest. Treatment with DMSO or Tunicamycin (Tm, 1 µg/mL, inhibitor of upstream oligosaccharide synthesis) were used as controls. DMSO treated lysates were split in half, with one half treated with PNGase F to remove *N*-linked glycans and induce a gel shift on glycosylated proteins. Probing for ERDJ3 (a glycosylated ER chaperone) was used to measure changes in glycosylation. GAPDH blotting used as loading control. The 5 µM dosage was chosen for subsequent infection experiments as it resulted in partial glycosylation inhibition while not completely suppressing MHV infection.
B. Western blots of DBT lysates treated with siRNA (40 h, Scramble, siCeacam1, or siMlec), then treated with NGI-1 (5 µM) or DMSO and infected with MHV-FFL2 (MOI 1, 10 hpi). Plain cells were untreated, uninfected to serve as background luminescence control. Lysates were probed for MLEC to verify knockdown, ERDJ3 to verify glycosylation inhibition, and GAPDH as loading control.

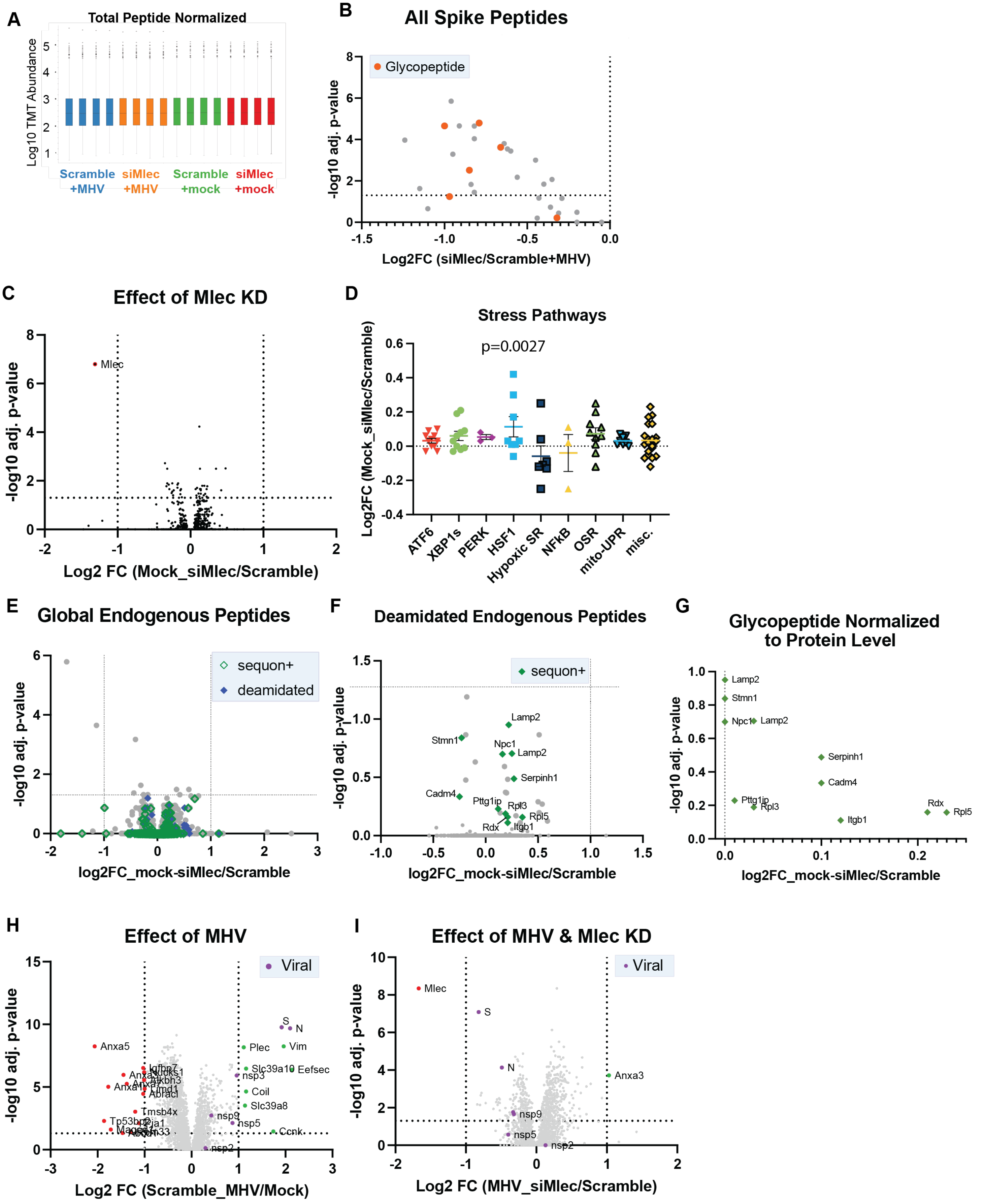

A
Total Peptide Normalized
Log10 TMT Abundance
Scramble +MHV
siMlec +MHV
Scramble +mock
siMlec +mock
B
All Spike Peptides
Glycopeptide
-log10 adj. p-value
Log2FC (siMlec/Scramble+MHV)
C
Effect of Mlec KD
Mlec
-log10 adj. p-value
Log2 FC (Mock_siMlec/Scramble)
D
Stress Pathways
p=0.0027
Log2FC (Mock_siMlec/Scramble)
ATF6
XBP1s
PERK
HSF1
Hypoxic SR
NFkB
OSR
mito-UPR
misc.
E
Global Endogenous Peptides
sequon+
deamidated
-log10 adj. p-value
log2FC_mock-siMlec/Scramble
F
Deamidated Endogenous Peptides
sequon+
Lamp2
Stmn1
Npc1
Lamp2
Serpinh1
Cadm4
Pttg1ip
Rpl3
Rpl5
Rdx
Itgb1
-log10 adj. p-value
log2FC_mock-siMlec/Scramble
G
Glycopeptide Normalized to Protein Level
Lamp2
Stmn1
Npc1
Lamp2
Serpinh1
Cadm4
Pttg1ip
Rpl3
Rdx
Rpl5
Itgb1
-log10 adj. p-value
log2FC_mock-siMlec/Scramble
H
Effect of MHV
Viral
S
N
Anxa5
Plec
Vim
Slc39a10
Eefsec
nsp3
Coil
Slc39a8
nsp9
nsp5
Ccnk
nsp2
Tmsb4x
-log10 adj. p-value
Log2 FC (Scramble_MHV/Mock)
I
Effect of MHV & Mlec KD
Viral
Mlec
S
N
Anxa3
nsp9
nsp5
nsp2
-log10 adj. p-value
Log2 FC (MHV_siMlec/Scramble)

**Figure S15. Glycoproteomics metrics for MHV infection.**

A. TMT channel abundance normalized to total peptide abundance. n = 4/condition, 1 MS run
B. Volcano plot showing the effect of Mlec-KD on Spike peptides in MHV infection (MHV-siMlec/MHV-Scramble). Glycopeptides are highlighted (orange). ANOVA test with multiple testing corrections for significance.
C. Volcano plot showing the effect of Mlec KD on the cellular proteome in uninfected DBT cells (Mock-siMlec/Mock-Scramble). Dotted lines indicate log2 FC < -1 or > 1 and adj. p-value <0.05. ANOVA test with multiple testing corrections for significance. Downregulated (red), upregulated (green), and viral (purple) proteins are annotated.
D. Protein marker levels of cellular stress pathways with Mlec KD in uninfected DBT cells (Mock-siMlec/Mock-Scramble), based on previously defined pathway gene sets(Davies et al., 2023; Grandjean et al., 2019; Shoulders et al., 2013). Dots represent individual proteins associated with each pathway. One-way ANOVA with Benjamini, Krieger and Yekutieli multiple testing corrections to test for significance, with p<0.05 annotated.
E. Volcano plot of global endogenous peptides with Mlec KD in uninfected DBT cells. Peptides that contain an *N*-glycan sequon (N-X-S/T) and/or are deamidated are annotated. Dotted lines indicate log2 FC < -1 or > 1 and adj. p-value <0.05.
F. Volcano plot of deamidated endogenous peptides as in (E). Peptides that contain an *N*-glycan sequon are annotated and labeled.
G. Glycopeptides from (F) normalized to the protein abundance of the associated protein.
H. Same as in (C), but showing the effect of MHV infection on Scramble-treated DBT cells (MHV-Scramble/Mock-Scramble).
I. Same as in (C), but showing the effect of Mlec-KD on MHV infection (MHV-siMlec/Mock-Scramble).

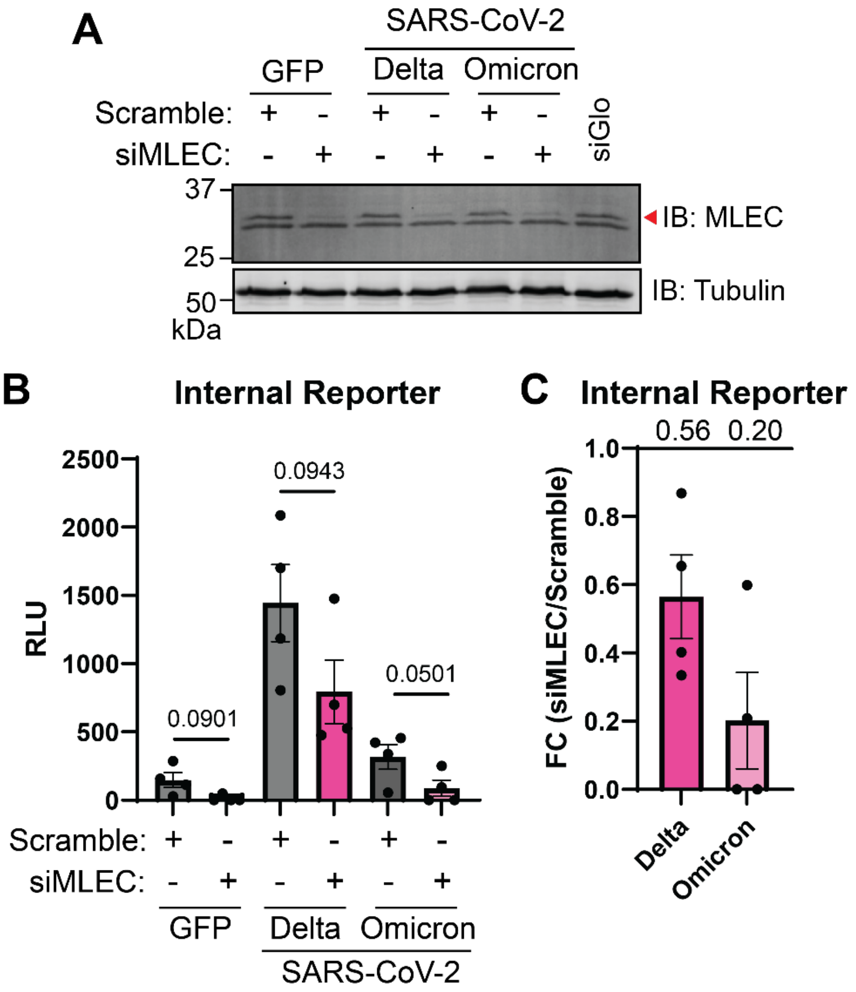

**Figure S16. Effect of MLEC knockdown on SARS-CoV-2 replicon internal reporter.**

A. HEK293T cells were transfected with siMLEC siRNA and SARS-CoV-2 replicons (Delta or Omicron), with Scramble siRNA, siGlo siRNA, and GFP transfection used as controls. After 72 h, cells were harvested and lysed in RIPA buffer with protease inhibitor. Lysates were analyzed by Western blot and probed for MLEC (top band, indicated with red arrow) and Tubulin (as a loading control). A representative Western blot is shown. n = 4
B. Lysates from (A) were assayed with a nanoGlo luciferase kit to measure levels of the replicon reporter that were not secreted. The total levels of signal were drastically lower than that of secreted reporter, indicating that the majority of reporter is secreted; however, the overall pattern of reduction of replicon with MLEC knockdown remains consistent. n = 4, Student's t-test for significance, with $p < 0.05$ considered significant. Bars represent mean values, ± SEM.
C. Data from (B) presented as a fold change comparison of siMLEC vs. Scramble treatment with respective SARS-CoV-2 replicon. Mean fold change value is annotated above the graph (0.56 and 0.20 respectively). Bars represent mean values, ± SEM.
